# The *Escherichia coli* O157:H7 Carbon Starvation-Inducible Lipoprotein Slp Contributes to Initial Adherence *In Vitro* via the Human Polymeric Immunoglobulin Receptor

**DOI:** 10.1101/429480

**Authors:** Christine Fedorchuk, Indira T. Kudva, Subhashinie Kariyawasam

## Abstract

*Escherichia coli* O157:H7 is the most well-studied serotype of enterohemorrhagic *E. coli* (EHEC) class of *E. coli* intestinal pathogens, and is responsible for many outbreaks of serious food-borne illness worldwide each year. Adherence mechanisms are a critical component of its pathogenesis, persistence in natural reservoirs, and environmental contamination. *E. coli* O157:H7 has a highly effective virulence operon, the Locus of Enterocyte Effacement (LEE), and its encoded intimate adherence mechanism is well characterized. However, factors involved in the preceding initial attachment are not well understood. In this study, we propose a mechanism of initial adherence used by *E. coli* O157:H7 *in vitro*. We describe a bacterial protein not previously reported to be involved in adherence, Slp, and its interactions with the human host protein polymeric immunoglobulin receptor (pIgR). The human pIgR has previously been shown to act as an adherence receptor for some mucosal pathogens, and is highly expressed in the intestine. Following observation of significant colocalization between *E. coli* O157:H7 and pIgR location on Caco-2 cells, a co-immunoprecipitation (Co-IP) assay using a human recombinant Fc-tagged pIgR protein led to the identification of this protein. Disruption of Slp expression in *E. coli* O157:H7, through deletion of its encoding gene *slp*, produced a significant adherence deficiency to Caco-2 cells at early time points associated with initial adherence. Plasmid complementation of *slp* fully restored the wild-type phenotype. Furthermore, immunofluorescence microscopy revealed evidence that this interaction is specific to the pathogenic strains of *E. coli* tested, and not the nonpathogenic strain E. coli K12. Additionally, deletion of *slp* resulted in the absence of the corresponding protein band in further Co-IP assays, while the plasmid-encoded *slp* complementation of the deletion mutant strain restored the wild-type pattern. These data support the proposal that Slp directly contributes to initial adherence, with the pIgR protein as its proposed receptor.

**Author summary:** *Escherichia coli* O157:H7 and other enterohemorrhagic *E. coli* (EHEC) are responsible for tens of thousands of cases of food-borne illness in the United States each year. *E. coli* O157:H7 has a particularly effective intimate adherence mechanism. However, the mechanisms of initial adherence, which facilitate attachment and virulence prior to the engagement of intimate adherence, are not well understood. In this study, we describe an initial adherence interaction between the *E. coli* O157:H7 Slp and the human polymeric immunoglobulin receptor (pIgR) expressed by the human colonic epithelial cell line Caco-2. The relationship was first demonstrated as a significant colocalization between the locations of *E. coli* O157:H7 bacterial cells and pIgR protein using immunofluorescence microscopy. The *E. coli* O157:H7 Slp protein was identified, and disruption of the *slp* gene resulted in a severe adherence deficiency to Caco-2 cells during initial adherence. This effect was reversed upon complementation of the Δ*slp* strain with a plasmid-encoded *slp* gene, and the constitutive over-expression of *slp* resulted in hyper-adherence exceeding that of the wild-type *E. coli* O157:H7. These data support the proposition that Slp directly contributes to initial adherence, with the pIgR protein as its proposed receptor.

## Introduction

It has been estimated that the most common enterohemorrhagic *E. coli* (EHEC) serotype, *E. coli* O157:H7, causes more than 70,000 cases of food-borne illnesses in the United States each year, with an additional 35,000 infections caused by non-O157 EHEC serotypes (1). EHEC infections typically result in abdominal cramping, watery or bloody diarrhea, and are also sometimes accompanied by fever, nausea, and vomiting (2). Among these cases, a subset develop a severe complication known as hemolytic uremic syndrome (HUS), which is primarily observed in infected children under the age of five years (3). In the U.S., *E. coli* O157:H7 outbreaks occurring between 1982 and 2002 had highly variable rates of HUS ranging from less than 1% to greater than 15% (4).

*E. coli* O157:H7 utilizes a fairly well-characterized set of virulence factors to colonize the human gastrointestinal (GI) tract during infection, including the robust intimate adherence mechanism carried out by factors encoded in the Locus of Enterocyte Effacement (LEE) pathogenicity island (5). The three main factors required for intimate adherence are (i) intimin, (ii) Tir, and (iii) type three secretion system (T3SS); which are the bacterial adhesin, translocated intimin receptor, and secretion apparatus, respectively (6). Upon infiltration of the mucin layer in the colon, the bacteria require an initial attachment in order to stay stably adjacent to the intestinal epithelial cell (IEC) for long enough to have the opportunity to engage its intimate adherence mechanism before being cleared from the digestive tract (7).

Several adhesins putatively involved in initial adherence have been reported, though the research has not yet reached a consensus. For example, long polar fimbriae (LPF) have been described as initial adhesins in *E. coli* O157:H7, but with some conflicting reports. LPF is a fimbrial adhesin with two loci, *lpf1* and *lpf2*, with homologs in *Salmonella* (8). Recently, LPF has been described in the role of translocation of EHEC across human M cells in *in vitro* human organ culture simulated conditions, but *lpf* deletion mutant strains of *E. coli* O157:H7 did not show any significant effects on adherence to Caco-2 cells after three hours post-infection (9). One study showed that a non-fimbriated *E. coli* K12 strain expressing the *lpf* operon demonstrated some increased adherence to MDBK and HeLa cells after six hours of infection, and single deletion mutations in *lpf1 or lpf2* showed a modest reduction in adherence under the same conditions (8). However, expression of *lpf2* in *E. coli* K12 was shown to result in reduced adherence to Caco-2 cells, so the exact role of LPF in initial adherence remains unclear (10). The EHEC factor for Adherence (Efa) has also been investigated as an initial adhesin in enteropathogenic (EPEC) strains, and *E. coli* O157:H7 contains a truncated version of *efa* (11). Insertion mutations disrupting the expression of *efa1* in EHEC showed reduced agglutination with human erythrocytes and autoagglutination with other EHEC cells, in addition to a significant reduction in adherence to Chinese hamster ovary (CHO) cells and a HeLa-derived cell line. However, the coinciding reduction in the secretion of the LEE-encoded effector EspA suggests that this effect may be more involved in intimate adherence than initial attachment (12) (13). Similarly, the effector ToxB found on the *E. coli* O157:H7 pO157 plasmid shares some sequence homology with Efa and has been shown to contribute to adherence. Its mechanism is likely to be involved with promotion of the LEE-encoded effector proteins EspA, EspB, and Tir and therefore is also unlikely to be involved in initial adherence (14). Flagella (H antigens) are well characterized for their role in motility, but have also been implicated in adherence in the bovine host. Flagella in *E. coli* O157:H7 were able to bind bovine intestinal mucus, while H negative mutant strains showed reduced adherence to bovine intestinal tissue explants; though other studies have reported H negative strains as being able to colonize young calves (6) (15). Although flagella have been shown to impact EPEC adherence *in vitro*, any role in EHEC or with human IECs remains unknown (16). Many other proteins have been identified as being of interest in the study of adherence, but roles in initial adherence are either unclear, not studied in human cells, or do not have enough data to be conclusive (7).

Under normal conditions, the human polymeric immunoglobulin receptor (pIgR) system plays a critical role in innate mucosal immunity (17). The human pIgR is a glycoprotein varying in size from 80–120 kDa depending on the level of glycosylation, and primarily functions as a transport for dimeric immunoglobulin A (dIgA) into the intestinal lumen (18). In the lumen, the transported pIgR-dIgA complex is cleaved from the IEC surface and termed secretory IgA (sIgA), where it plays an important role in maintaining the balance between immune function and microflora homeostasis (19) (20). In addition, pIgR lacking dIgA can also be transported. The unbound pIgR is also cleaved and released (free secretory component or SC), which also serves as part of innate immunity (21) (22). In spite of its importance in immunity, the human pIgR is also known to be hijacked by pathogens and used as a route for adherence and invasion. For example, the human pIgR is known to contribute to the adherence and invasion of *Streptococcus pneumoniae*, and the invasion and intercellular spread of Epstein-Barr virus during infection (17) (23). Although *S. pneumoniae* is a Gram-positive pathogen, it is a pathogen that targets mucosal epithelial cells, as does EHEC (24) (25) (26). Additionally, a study in 2007 found that *pigr expression at six hours post-infection in bovine hosts was upregulated when* experimentally infected with *E. coli* O157:H7 and a non-O157 EHEC strain (1.93 and 3.51 fold-change (FC), respectively); but not in response to non-colonizing control strains (27). Although this effect on gene expression may have been due to the bovine host immune response to EHEC colonization, as a mucosal epithelium colonizing pathogen EHEC also has the potential to use the pIgR during adherence to IECs during infection.

In this study, we describe the interaction between the *E. coli* O157:H7 outer membrane protein Slp (carbon starvation-inducible lipoprotein) and the pIgR, and its role in initial adherence to Caco-2 cells *in vitro*. Utilizing immunofluorescent microscopy to investigate the possibility, we demonstrated that during initial adherence, *E. coli* O157:H7 had a significant correlation with its location and the location of pIgR protein on the Caco-2 cell surface. The colocalization of *E. coli* cells with pIgR protein was statistically significant with *E. coli* O157:H7 (O157:H7-WT), but not with the nonpathogenic *E. coli* strain K12. Additionally, several other non-O157 pathogenic *E. coli* strains showed significant colocalization patterns. These observations were further investigated using a commercially available recombinant Fc-tagged human pIgR protein (pIgR-Fc) in a co-immunoprecipitation (Co-IP) assay, leading to the identification of Slp as the interacting protein involved in the colocalization phenotype observed in the pathogenic strains of *E. coli* tested. A deletion mutation of *E. coli* O157:H7 lacking the *slp* gene (O157:H7-Δ*slp*) resulted in an elimination of the colocalization patterns observed with immunofluorescent microscopy, the absence of the ∼20 kDa band corresponding to Slp following Co-IP, and a significant adherence deficiency to Caco-2 cells. Furthermore, the O157:H7-Δ*slp* strain was complemented *in trans* with the *slp* gene (O157:H7-p::*slp*), which restored the wild-type phenotypes seen in colocalization, Co-IP, and adherence to Caco-2 cells *in vitro*. Taken together, these results support the direct interaction between the *E. coli O157:H7* Slp and the host pIgR and its contribution to initial adherence.

## Results

### Initial adherence timeline of *E. coli* O157:H7 during adherence to Caco-2 cells

In order to effectively study the roles of different factors during attachment, a specific timeline was required to define initial adherence in this model. Using the expression of LEE-encoded genes as a benchmark for the beginning of the intimate adherence stage of attachment, significant LEE upregulation has been cited as taking place between two and six hours post-infection *in vitro*. When adhered to Caco-2 cells, *E. coli* O157:H7 Sakai demonstrated increased production of EspA, EspB, and Tir protein production after 4.5 hours post-infection (28). During adherence to HeLa cells, LEE1 in *E. coli* O157:H7 Sakai has been shown to be activated by four hours post-infection (29) (30); while *eae* expression in an undefined *E. coli* O157:H7 strain demonstrated upregulation as early as two hours after infection (31). Other studies of EPEC have shown LEE promoter or LEE-encoded gene upregulation between four and six hours in several different epithelial cell lines, but the differences in the LEE operon between EPEC and EHEC strains make the relevance to *E. coli* O157:H7 unclear (32) (5). Following previous work, under the conditions of this study the intimin-encoding *eae* gene was used as a benchmark for the onset of intimate adherence. Compared to a non-adhered control culture to isolate the gene expression profiles of only initially attached bacterial cells, quantitative PCR (qPCR) was used to calculate FC of *eae* expression in adhered *E. coli O157:H7 bacteria was calculated using the relative* expression ratio method for comparative Ct analysis (33). The expression of *eae surpassed the* threshold for significance (FC ≥ 2) after four hours and increased thereafter, consistent with previous literature (Fig 1). Thus, initial adherence in this model was defined as taking place from zero to three hours post-infection, when *eae* was not significantly upregulated.

**Fig 1.**
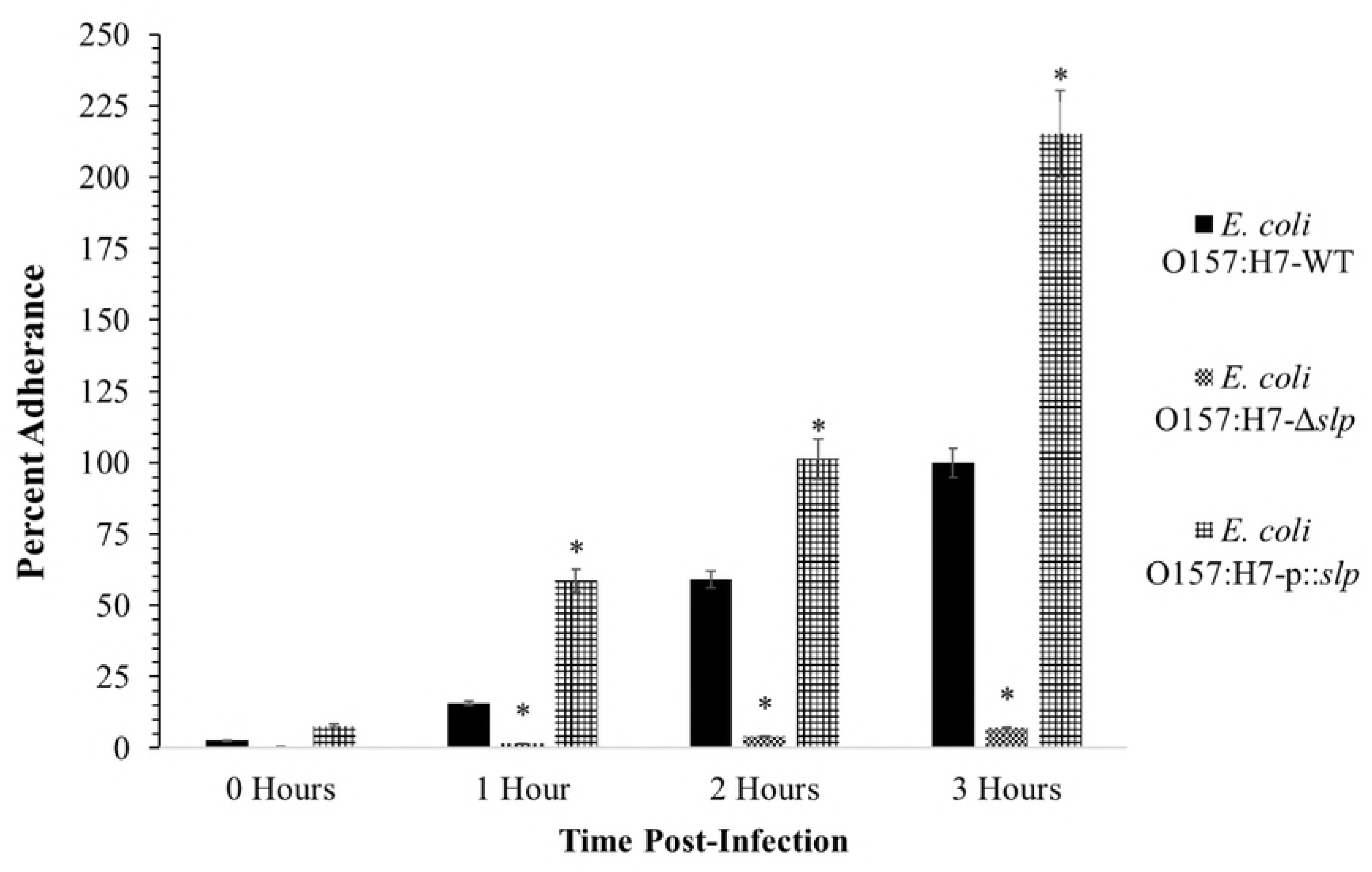
Relative expression of *eae* during *E. coli* O157:H7 adherence to Caco-2 cells and the initial adherence timeline of *E. coli* O157:H7 *in vitro*. Relative expression of *eae is shown as the fold-change over six hours post-infection, as* compared to *E. coli* O157:H7 grown in cell culture media alone. Significant upregulation is defined as FC ≥ 2.0. Threshold for significance is shown at FC = 2.0, and initial adherence timeline shown in the boxed area.

### Colocalization and covariance of pIgR with *E. coli*

To investigate the possibility of a pIgR adherence mechanism in *E. coli O157:H7 initial* adherence, immunofluorescence was used to assess colocalization between the location of pIgR protein and locations of adhered *E. coli* bacteria and quantify any correlation using covariance analysis (Fig 2). The covariance, shown as R, attested that there was no significant correlation between the locations of *E. coli* K12 bacteria and the pIgR protein; while O157:H7-WT did exhibit a significant correlation (Fig 2B-C). This relationship is further detailed when observed over time, where O157:H7-WT showed an initial peak of covariance (R > 0.7) at one hour post-infection, and decreased over two and three hours (R > 0.5 and 0.4, respectively). This pattern of early peak followed by steady decrease was consistent with the predicted pattern of an adhesin active during initial adherence, as its activity would be less necessary as the intimate adherence factors assumed adherence function at later timepoints. No such pattern was observed in *E. coli* K12. Several other extra-intestinal (ExPEC) pathogenic strains of *E. coli* were also tested and showed significant covariance at two hours post-infection, indicating the possibility that an adherence mechanism is not limited to only *E. coli* O157:H7 (Fig 2A-B).

**Fig 2.**
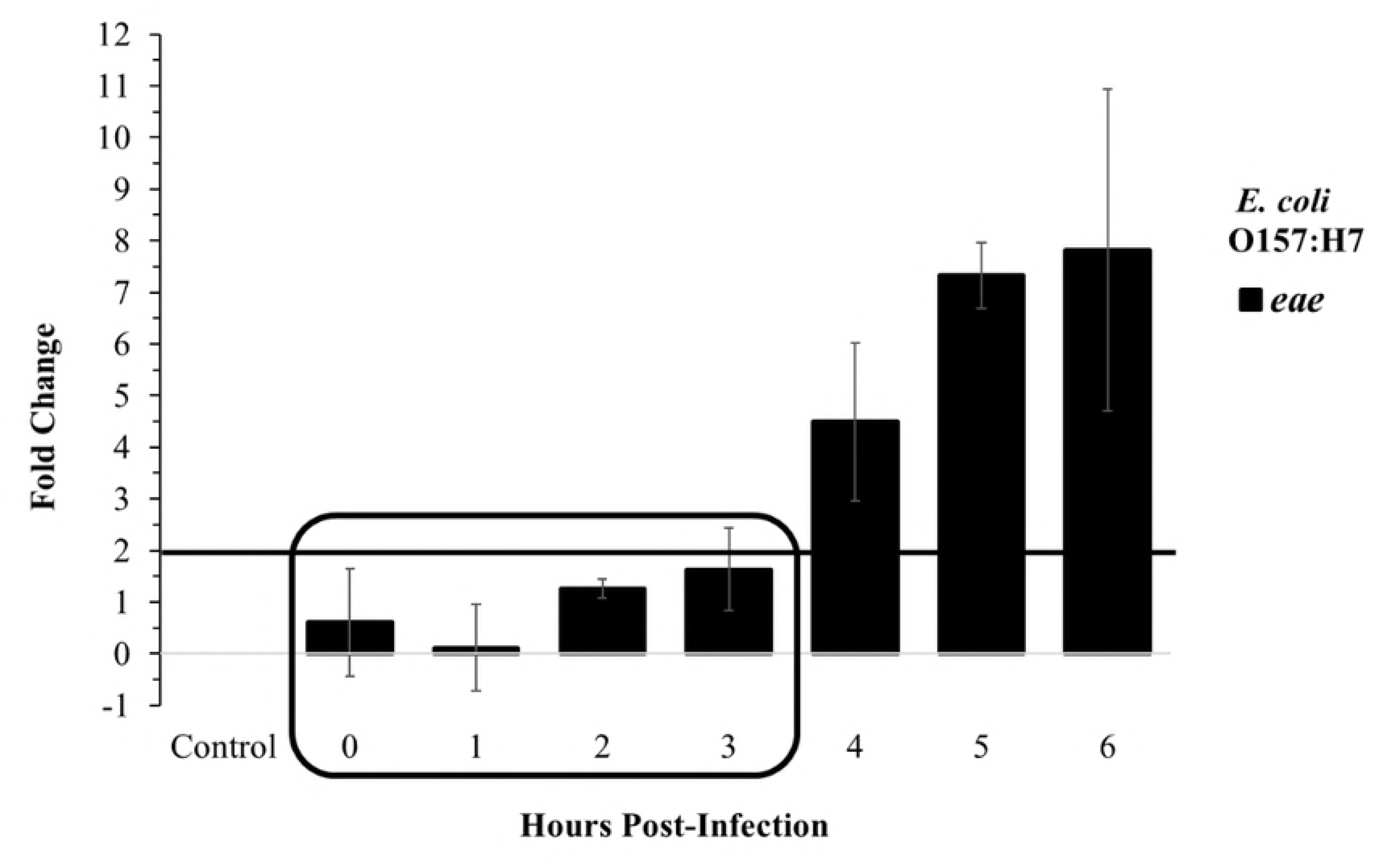
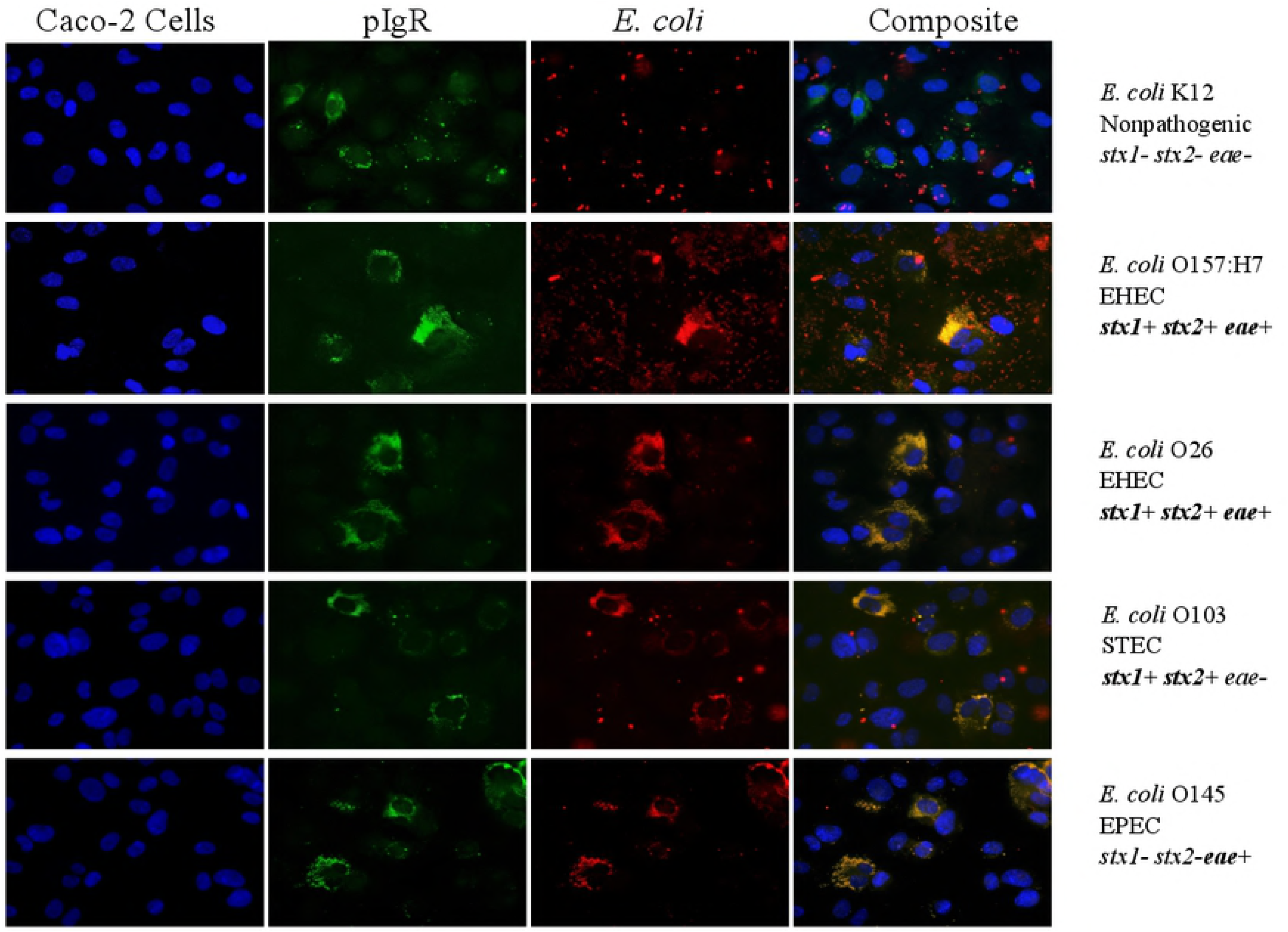
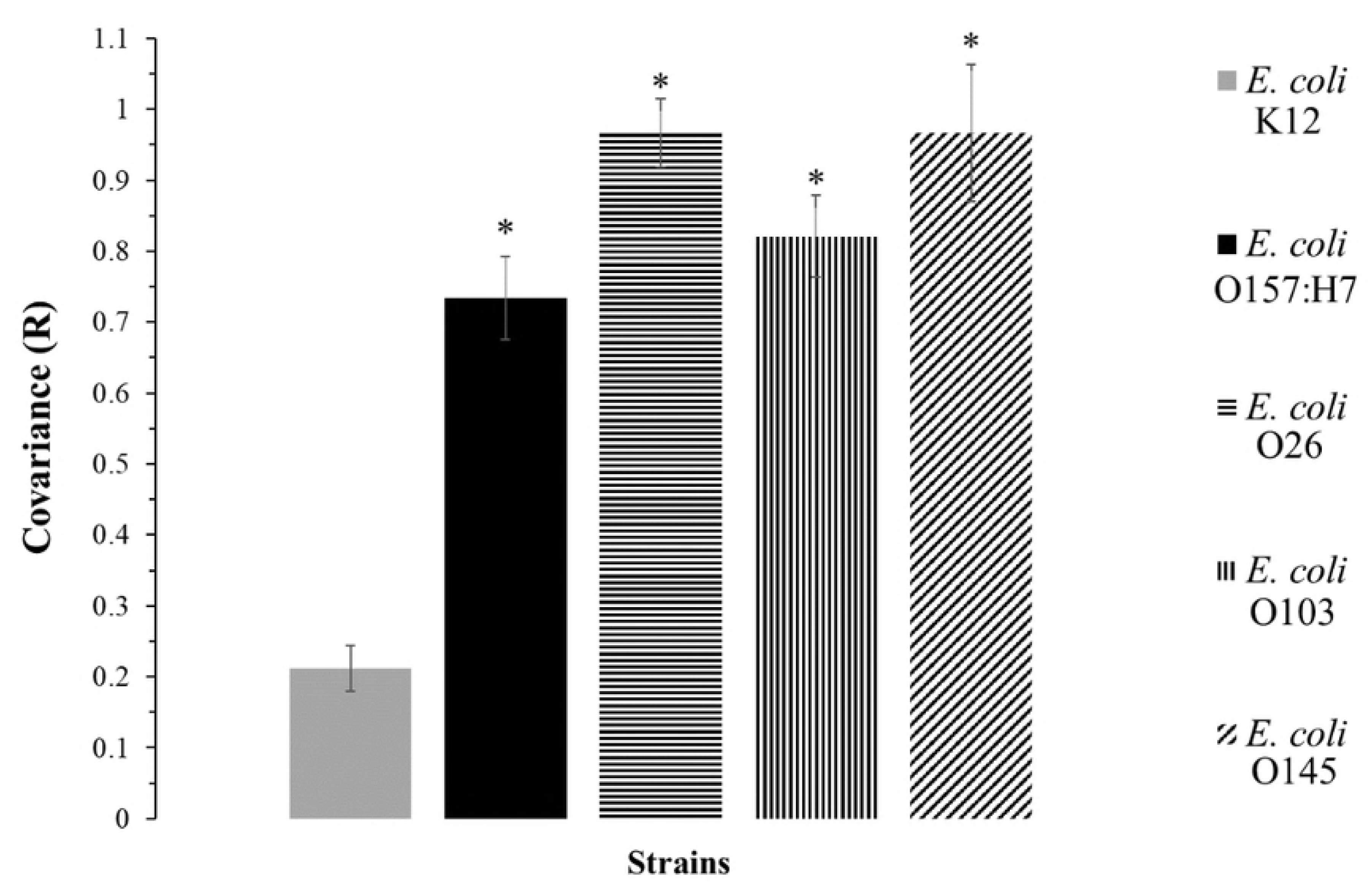
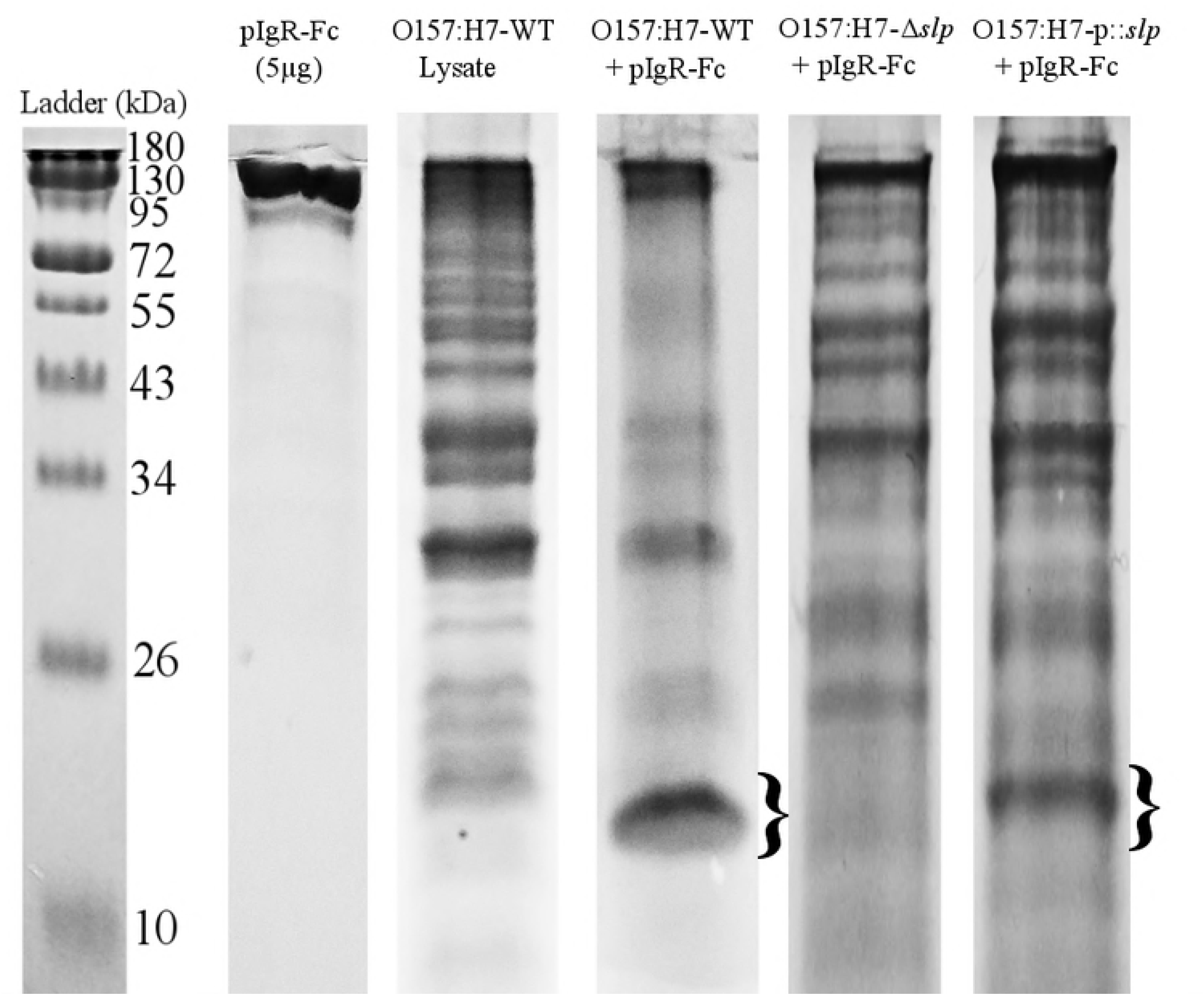
Colocalization and covariance of pIgR with *E. coli* strains during initial adherence. (A) Colocalization of pIgR with *E. coli* strains. Fluorescent signals of Caco-2 nuclei, Hoechst stain with emission at 497 nm, shown in blue; fluorescent signals of *E. coli*, DyLight 594 with emission at 594 nm, shown in red; fluorescent signals of pIgR protein, DyLight 488 with emission at 488 nm, shown in green; covariance of pIgR protein with *E. coli* K12 and O157:H7-WT (B) Covariance of pIgR with *E. coli* strains and (C) pIgR with *E. coli* K12 and O157:H7-WT over time. P-values were calculated using the mean PCC (R value) for the Student’s T-test and p ≤ 0.05 was considered significant.

### Protein-protein interactions between *E. coli* O157:H7 and the pIgR

To demonstrate a direct protein-protein interaction between *E. coli* O157:H7 and the pIgR, a co-immunoprecipitation (Co-IP) assay was done using a C-terminal Fc tagged human recombinant pIgR protein (pIgR-Fc) and O157:H7-WT proteins (whole-cell lysate was used to avoid the accidental exclusion of any proteins). Recovered proteins were run in a reducing SDS-PAGE gel (Fig 3), and bands of interest were later identified through LC MS/MS analysis (Table 1).

**Table 1.** Protein identification using LC MS/MS. Summary of the outer membrane proteins found to be possible identifications of the protein recovered from Co-IP. Proteins are listed in order of decreasing sequence coverage, and proteins that were identified in three of three LC MS/MS analyses are noted.

| Protein ID | Protein Description | Reference Sequence and Accession | Average % Coverage | # AAs | kDa |
| --- | --- | --- | --- | --- | --- |
| *Slp | Outer membrane protein Slp | *Escherichia coli* O157:H7 Str. EDL933 AAG58638.1 | 54.9 | 199 | 22.2 |
| *Pal | Peptidoglycan-associated lipoprotein | Multispecies WP_001295306.1 | 53.8 | 173 | 18.8 |
| *OmpW | Outer membrane protein W | Multispecies WP_000737224.1 | 49.8 | 212 | 22.9 |
| *OmpX | Outer membrane protein X | Multispecies WP_001295296.1 | 44.7 | 171 | 18.6 |
| OmpA | Outer membrane protein A precursor | *Escherichia coli* O157:H7 Str. EDL933 AIG67419.1 | 22.2 | 354 | 38.1 |
| OmpC | Outer membrane protein C precursor | *Escherichia coli* WP_000865552.1 | 15.4 | 367 | 40.3 |

\* Proteins present in all three LC MS/MS analyses.

Lane D shows the presence of a concentrated band of approximately 20 kDa (noted as }) in the treated sample, not visible in the untreated control sample (Lane C), indicating the likelihood of a direct protein-protein interaction with the pIgR-Fc protein (Fig 3). There were several other observable bands in the treated sample between 26–43 kDa, but they corresponded in size to the most concentrated bands observed in the control, making it likely that they were non-specific and not indicative of protein binding.

**Fig 3.**
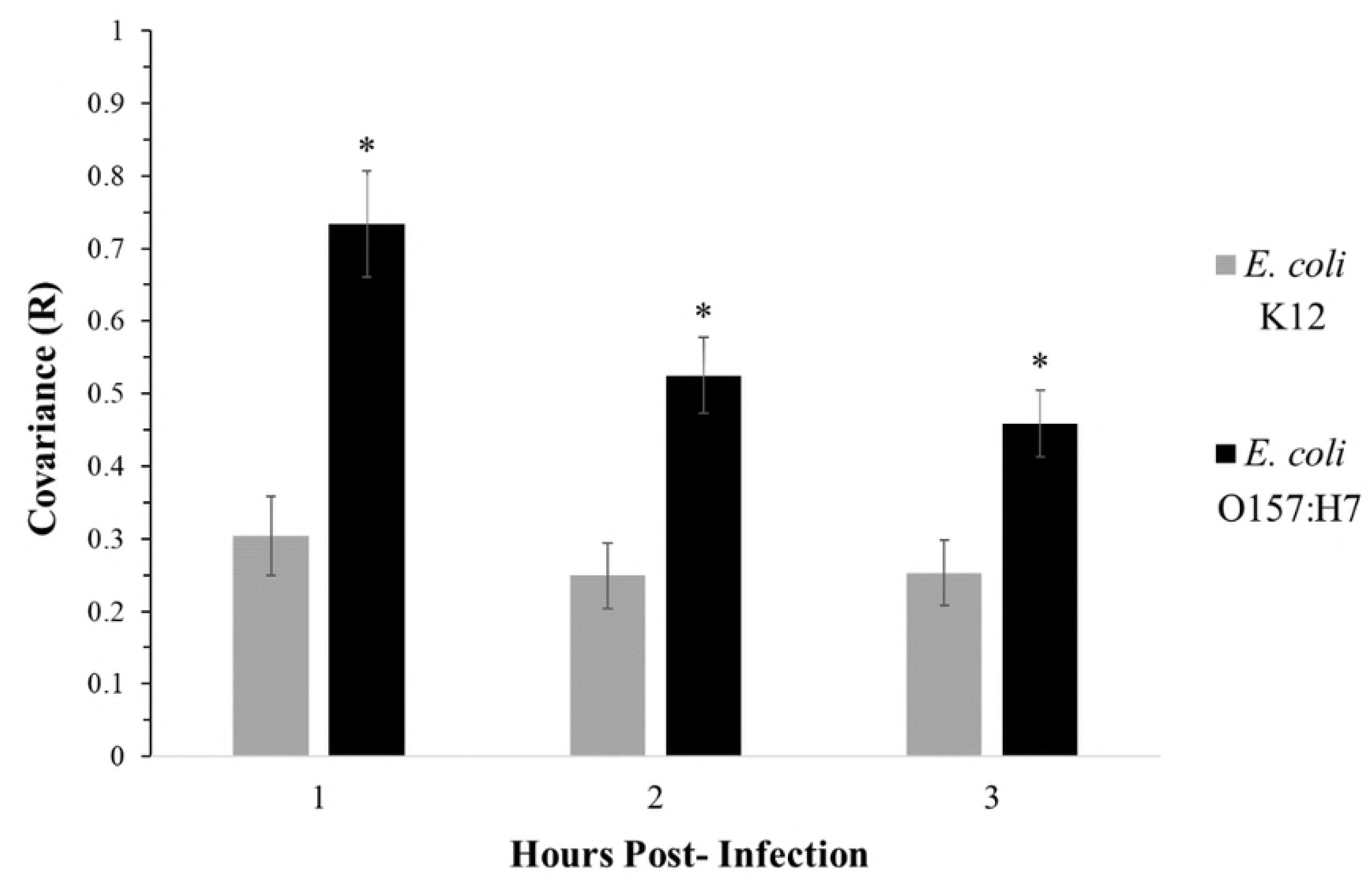
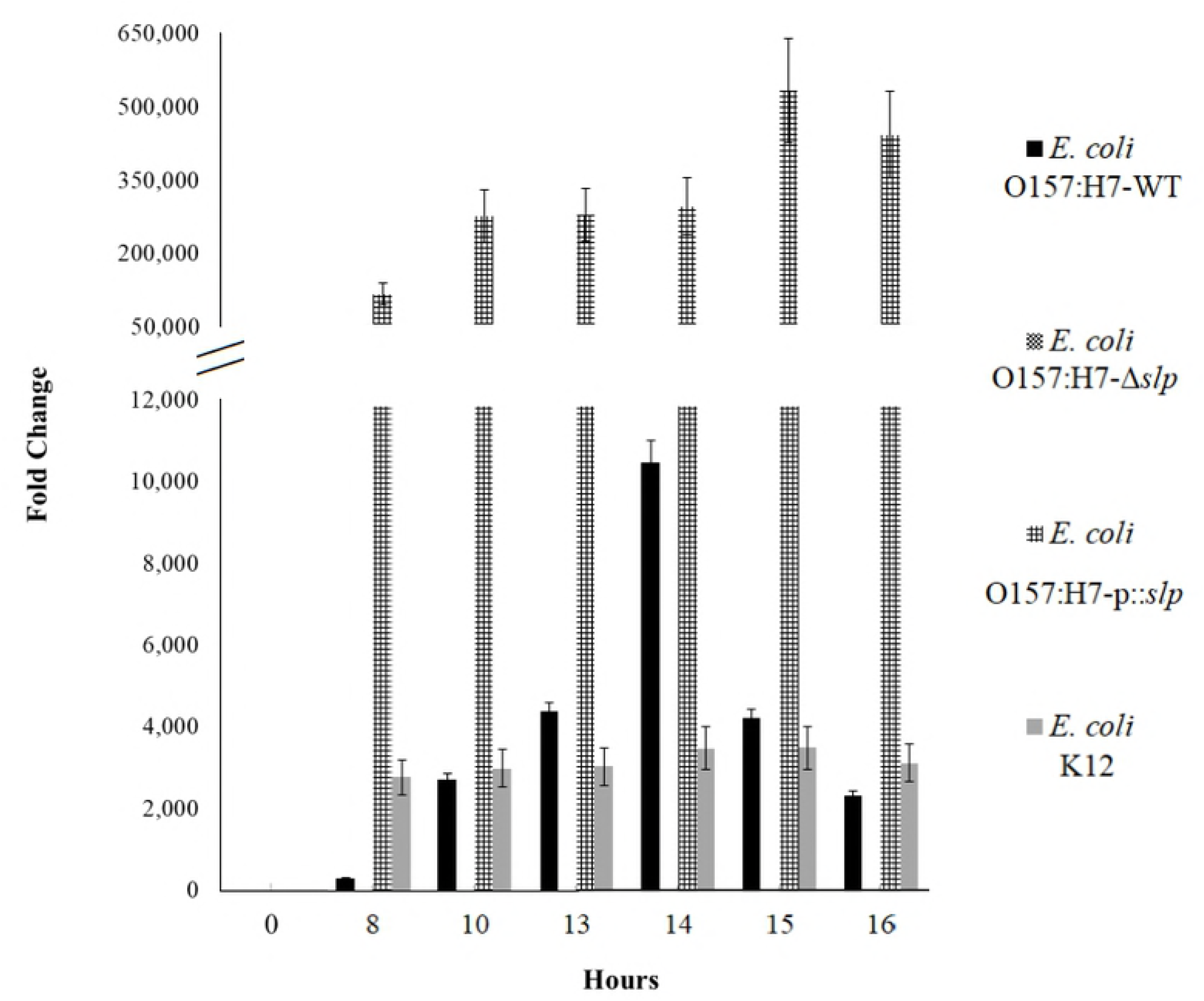
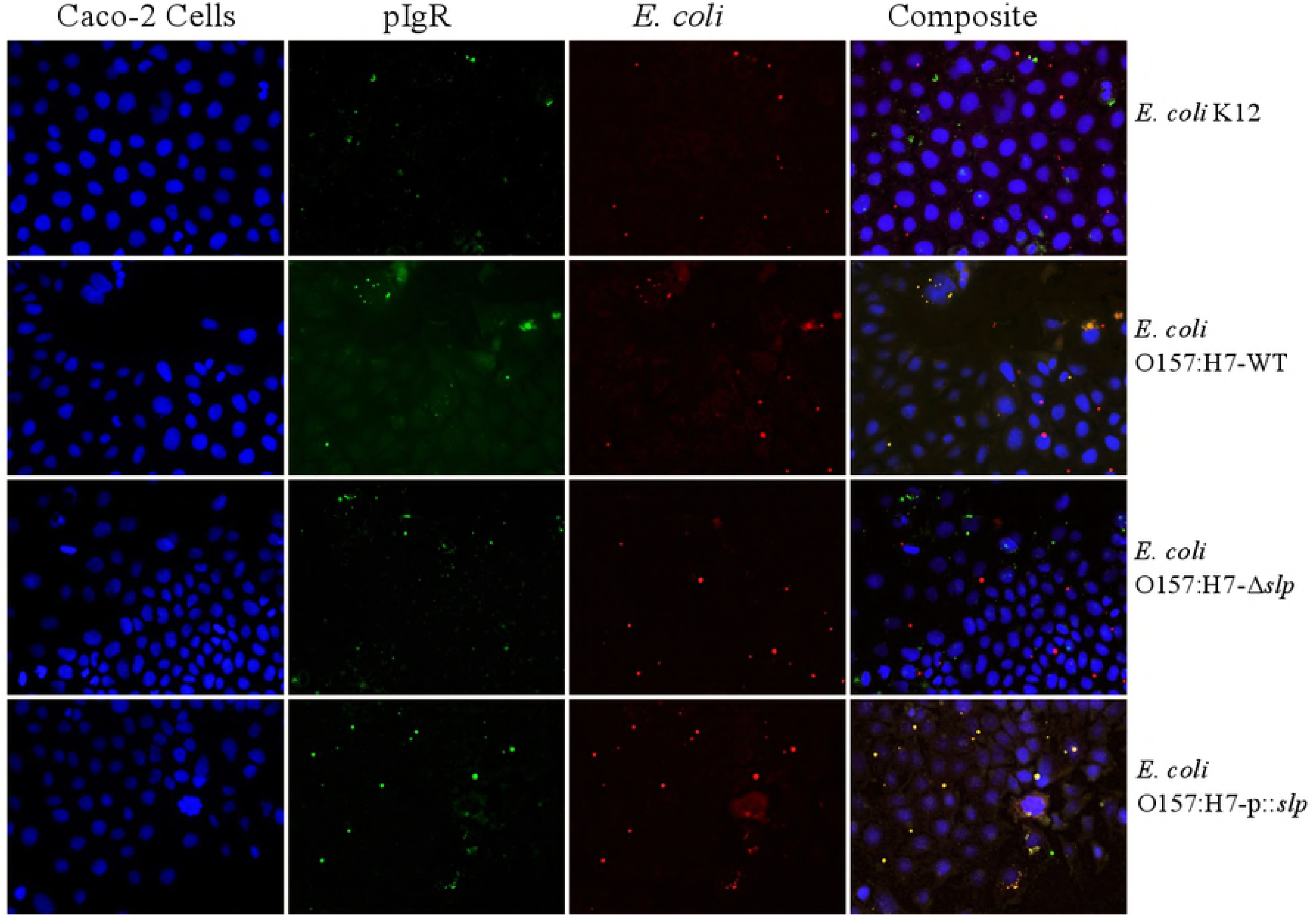
Co-immunoprecipitation of *E. coli* O157:H7 proteins with human recombinant Fc-tagged pIgR protein. SDS-PAGE showing the apparent sizes of proteins in the conditions used for Co-IP.

A direct relationship between the O157:H7-WT protein Slp and the pIgR-Fc protein was demonstrated through disruption of the *slp* gene. A deletion mutation of the *slp* gene (O157:H7-Δ*slp*) eliminated the band of interest recovered with the wild-type strain (O157:H7-WT) (Lane E), and the subsequent complementation of a plasmid-encoded *slp* gene (O157:H7-p::*slp*) restored the wild-type phenotype (Lane F) (Fig 3).

### Slp and initial adherence *in vitro*

Relative expression of the *slp* gene showed differing patterns between *E. coli* K12, O157:H7-WT, O157:H7-Δ*slp*, and O157:H7-p::*slp* as measured by qPCR (Fig 4A). The *slp* gene is reported to be transcribed at a low baseline level in most conditions and upregulated during entry of bacteria into stationary-phase (34). Over 16 hours of growth, O157:H7-WT *slp* expression showed a transient peak at 14 hours, corresponding to the entry of the culture into stationary-phase growth. *E. coli* K12 *slp* expression remained relatively stable and low throughout growth, and did not show any peaks or changes at any time point. The O157:H7-Δ*slp* strain did not show any detectable levels of gene expression, while the O157:H7-p::slp strain showed high levels of constitutive over-expression of *slp* at high levels at all timepoints, ranging from 10 to over 100 magnitude differences in FC as compared to *E. coli* K12 and O157:H7-WT.

**Fig 4.**
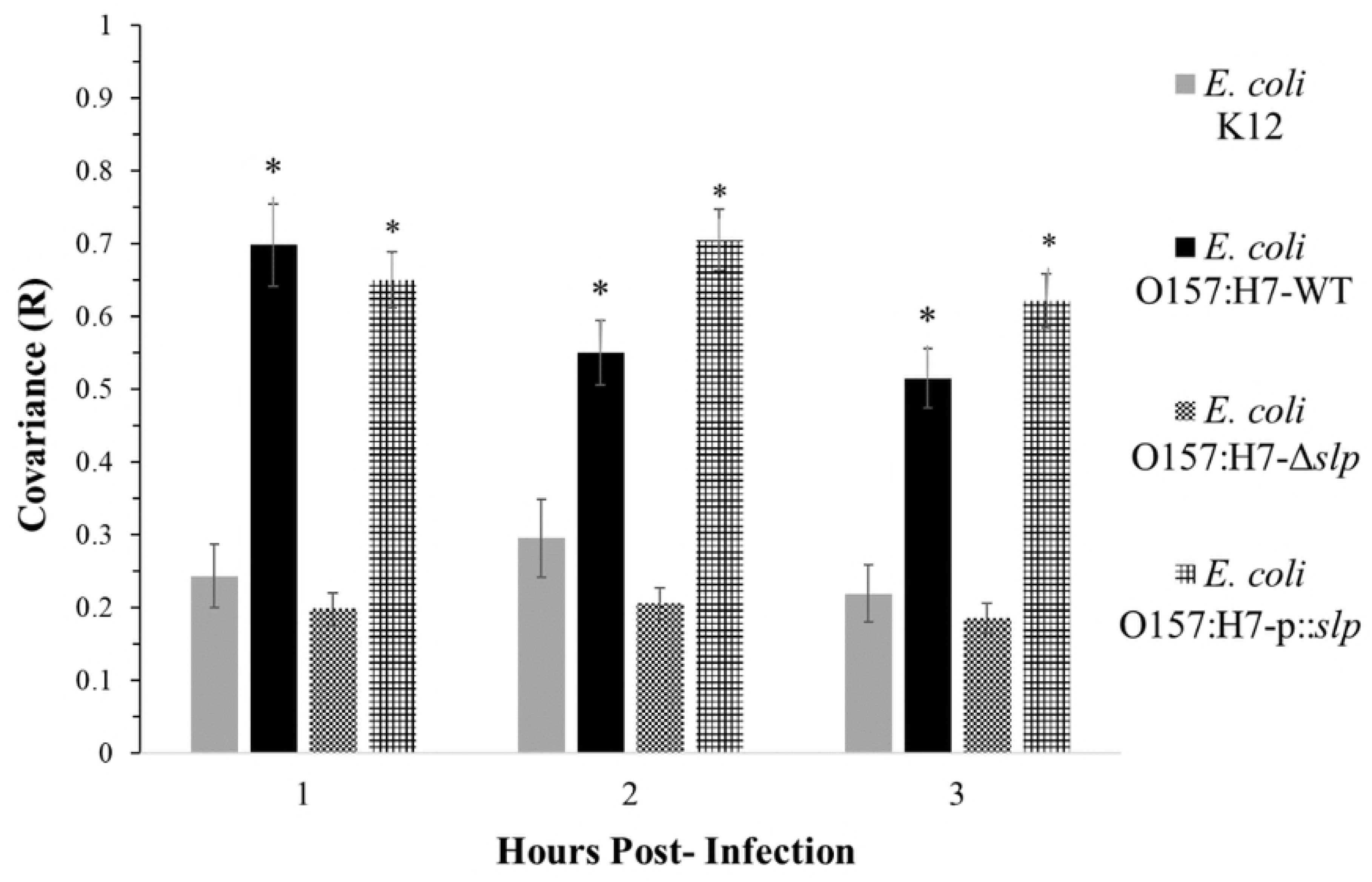
Relative *slp* gene expression, covariance, and adherence in *E. coli* strains. (A) Relative gene expression of *slp* in culture. (B) Quantitative adherence of *E. coli* strains to Caco-2 cells over time, relative to the maximum number of adhered wild-type bacteria (100% adherence is wild-type at three hours post-infection. (C) Colocalization and (D) covariance of pIgR with *E. coli* O157:H7-WT, O157:H7-Δ*slp*, and O157:H7-p::*slp*. P-values were calculated using the mean PCC (R value) for the Student’s T-test and p ≤0.05 was considered significant.

When measuring initial adherence to Caco-2 cells by quantifying the number of attached bacteria over time, the O157:H7-WT presented a pattern showing a steady increase of adhered bacterial cells over three hours (Fig 4B). These results were consistent with O157:H7-WT bacterial cells as they were steadily coming into contact with and attaching to the Caco-2 cells over time, as the adherence increased two- to three-fold per hour (approximately 20%, 55%, and 100% at 1, 2, and 3 hours post-infection, respectively). When compared to the wild-type, deletion of the *slp* gene resulted in a significant adherence deficiency to Caco-2 cells, with O157:H7-Δ*slp* adherence not reaching more than 10% at any time point. The O157:H7-Δ*slp* adherence increased very slightly over three hours (from <5% to <10% at 1 and 3 hours post-infection, respectively). Conversely, the O157:H7-p::slp showed a hyper-adherent pattern, surpassing wild-type levels at all timepoints. The O157:H7-p::*slp* adherence followed roughly the same pattern as seen in the wild-type, with adherence increasing two- to three-fold per hour (approximately 60%, 100%, and 225% at 1, 2, and 3 hours post-infection, respectively). At three hours, the number of adhered O157:H7-p::*slp* cells was more than double the wild-type.

Colocalization and covariance were measured in the O157:H7-Δ*slp* and O157:H7-p::slp strains over three hours of adherence to Caco-2 cells (Fig 4C-D). As in Fig 2, *E. coli* K12 did not show any statistically significant colocalization or any changes over time (R = 0.25, 0.28, and 0.22 at 1, 2, and 3 hours post-infection). The O157:H7-WT cells showed a significant colocalization at all timepoints, with the highest correlation (R = ∼0.7) seen at one hour post-infection, with a decrease to R = 0.55 and 0.50 at two and three hours respectively. The O157:H7-Δ*slp* strain demonstrated adherence levels comparable to *E. coli* K12, with significantly diminished colocalization and no changes over time (R = approximately 0.2 at all time points). The O157:H7-p::*slp* strain restored colocalization comparable to that of the wild-type strain, but colocalization did not decrease over time in the same manner (R = 0.65, 0.70, and 0.63 at 1, 2, and 3 hours post-infection, respectively).

## Discussion

Initial adherence of EHEC bacteria is a known step in pathogenesis- a loose attachment between the bacterium and the epithelial cell is required in order to have the opportunity to make an intimate attachment using T3SS effectors (35). The initial adherence in EHEC strains is not yet fully understood, and it is becoming clear that mechanisms vary between strains, hosts, and environmental signaling factors present (7). Studies examining initial adhesins have used a diverse array of host species as models with a variety of EHEC strains, which have shown that adherence mechanisms are not conserved. Even among EHEC strains with very similar virulence profiles, putative adhesins have inconsistent behavior between host type (7) (36) (37) (14). For example, in *E. coli* O157:H7, the flagellar antigen H7 has been implicated in bovine cell adherence using bovine intestinal tissue explants (38). Flagella in *E. coli* O157:H7 were able to bind bovine intestinal mucus, while H negative mutant strains showed reduced adherence to bovine intestinal tissue explants (6). However, purified H7 protein did not show any ability to bind to HeLa cells when other purified H proteins were able to bind in the same conditions (39). Here, using human colonic epithelial cells, we demonstrated a novel EHEC initial adherence mechanism *in vitro*, involving the *E. coli* O157:H7 outer membrane protein Slp and the human pIgR protein.

The role of the pIgR in pathogenesis is a known mechanism for other pathogens, but it had not been demonstrated in *E. coli* adherence. The pIgR is a major component of innate mucosal immunity, and its regulation is influenced by a variety of factors activated as part of the general inflammatory response to pathogens during colonization and infection (21). It was speculated that the pIgR behavior observed could have been affected by an immune response to the presence of EHEC; however, the effects of EHEC on the pIgR system do not explain the results reported. Like many enterics, *E. coli* is a Gram-negative organism, and contains LPS in its membrane (40). The LPS molecule is a potent activator (and the only known ligand) of Toll-like receptor 4 (TLR4), which causes an increased expression of the transcription factor NF-κB when activated (41). NF-κB is a highly effective activator of multiple proinflammatory signals, and among other effects it induces transcription of the *pigr* gene (42). Shiga toxins (Stx) are also capable of inducing expression of *pigr through an induction of proinflammatory cytokine IL-8* during ribotoxic stress response *in vitro* (43). However, in order to upregulate *pigr adequately to* effect attachment as early as zero to three hours post-infection, exposure of the host cell to LPS or Stx would have to be reasonably prompt, prolonged and stable; which is unlikely to occur in the variable intestinal environment (41). Additionally, there is no evidence to suggest that the pIgR has any mechanism of location specificity, and inflammatory signals only affect the rate at which pIgR reaches the membrane without any effect on its location (19) (44) (45).

In this study, we hypothesized that the pIgR may be involved in EHEC adherence, and determined the bacterial protein binding to the human pIgR protein using a Co-IP assay. The Co-Escherichia coli through interactionsIP produced a distinct band of approximately 20 kDa that was significantly more concentrated when *E. coli* O157:H7 proteins were incubated with pIgR-Fc than with *E. coli* O157:H7 proteins alone. Identification using LC MS/MS led to the *E. coli* O157:H7 outer membrane protein Slp, a 22 kDa lipoprotein primarily known to be expressed in *E. coli* during entry into stationary-phase and during carbon starvation (34). Its encoding gene (*slp*) expression is extremely low at every other growth stage, only being upregulated significantly during entry into stationary-phase growth or during carbon starvation. Its function is not completely clear, though it has generally been accepted that one of Slp’s probable functions is related to membrane stability during stationary-phase growth (34) (46). The O157:H7-Δ*slp* strain was observed to be slightly more easily lysed in water than the wild-type strain, which supports these findings. Bacterial lipoproteins are a broad range of proteins and have been known to be involved with many virulence functions, including adherence (47). Other genes with low levels of sequence homology have been identified in other Gram-negative bacteria or enteric pathogens, which suggests the possibility of a similar role among some limited classes of bacteria (46). The *slp* sequence is located on a genomic acid fitness island (AFI) along with several other defined and undefined genes and operons (48). The *slp* gene and AFI are present in all *E. coli*, but there are significant differences between strains. In *E. coli* K12, the AFI is approximately 14 kb and contains 12 protein-encoding genes, whereas the *E. coli O157:H7 AFI is 23 kb and contains 21* protein-encoding genes. Due to the insertion of an O-island sequence encoding a heme transport locus, the *E. coli* O157:H7 AFI contains nine extra genes involved in heme uptake and utilization (*chuA* genes) (48) (49) (50). The *slp* is located in a small operon and is transcribed along with the gene *yhiF* (which encodes a putative LuxR family regulator protein), but of the AFI proteins, Slp and YhiF are not well characterized (49) (51). Several studies have outlined different potential functions of these proteins and how they may function together or separately, but the data are not yet conclusive. A 2007 study proposed that Slp and YhiF function together to protect the *E. coli* cell from its own toxic metabolic products during organic acid metabolism (52). Another study demonstrated the antimicrobial effects of cranberry concentrate in ground beef via bacterial membrane damage, and reported the differential expression of several outer membrane protein-encoding genes, including *slp*. The *slp* gene was modestly downregulated in conditions that showed anti-microbial activity, indicating a possibility of membrane weakening through lack of Slp’s function in membrane stabilization (53). There has also been evidence of increased *slp* gene expression during biofilm formation, though the significance of these findings is unclear (54). It is notable that Slp contributes to initial adherence and is involved in acid stress response and resistance, because pH is a potent signaling factor in *E. coli* O157:H7, and resistance to acid stress is a major component of *E. coli* O157:H7 virulence (55). Acid stress resistance in *E. coli* O157:H7 is crucial to survival in the low pH environment of the stomach, and the most significant AR mechanism used is the glutamate-decarboxylase (Gad) system, which is also found in the AFI (49). The regulation of adherence and virulence genes affected by pH is notoriously complex and not well understood; particularly with the Gad system (56). However, it has been shown that low pH results in a gene expression profile fit for initial adherence (55). In response to low pH, Gad-related regulators will downregulate the expression of LEE-encoded effectors not required at that stage of early attachment (i.e. *eae and other intimate adherence* related genes), and upregulate genes related to adherence and motility (55) (57) (49) (58). GadE, the Gad system master regulator, is known to affect the expression of genes outside the AFI both directly and indirectly (50). An acid- or Gad system-influenced mechanism of expression would be beneficial for initial adherence factors, as it is a consistent and robust signal present prior to the environment where initial adhesins would be required for effective colonization (58). With many virulence factors converging to be influenced or controlled by these acid-regulated pathways, regulation of adherence factor expression in the same manner would be more reliable and efficient than having multiple independent mechanisms during pathogenesis (59). Additionally, elaborate regulatory mechanisms also provide insight into possible reasons why *E. coli* K12, despite having the slp gene and a similar AFI, does not show a colocalization phenotype with pIgR. While no work has yet been done specifically regarding adherence and AFI-encoded gene regulation in response to acid stress in *E. coli* K12, the differences in *slp* expression in culture over time provide evidence that *slp* expression is managed by a different mechanism in *E. coli* K12 than in *E. coli* O157:H7. A study from 2011 highlighted some significant differences between *E. coli* K12 and *E. coli* O157:H7 AFI gene expression-namely, that *E. coli* O157:H7 carries a mutation causing the *hdeB* gene from being expressed (48). HdeA and HdeB are AR chaperone proteins that play a critical role in acid tolerance in *E. coli* K12. Deletion mutations of *hdeA, hdeB*, or *hdeAB* will significantly hinder the ability of *E. coli* K12 to survive acid stress, but *E. coli* O157:H7 was almost completely unaffected by the deletion of these genes (48). These findings illustrate the presence of different AFI gene expression regulation mechanisms and implies that these difference may explain why an AFI-containing *E. coli* strain does not appear to utilize the Slp during adherence. Given the complexity and sensitivity of acid-responsive regulatory mechanisms, and the current gap in knowledge regarding the exact roles of Gad, AFI, and other regulatory genes during infection of the human host; further study is required to fully understand these systems and how their gene products, such as Slp, affect pathogenesis *in vivo*.

*E. coli* O157:H7 and other EHEC strains pose a large public health burden worldwide, and with the most serious complications and outcomes affecting children under the age of five years, the need for effective treatments or interventions is urgent (3). Interventions are aimed at preventing the transmission from the colonized bovine host at either the pre- or post-harvest stage, by treating or preventing EHEC colonization before slaughter, or reducing or eliminating contamination post-harvest (60). Results of interventions such as vaccines and post-harvest sanitation have produced limited effects (61) (62) (60) (63). Many of the EHEC virulence and adherence related genes are significantly affected by an acidic environment, and this convergence of regulatory systems involving adherence and virulence provides a potential opportunity for interventions or therapies to target these pathways and attenuate virulence or adherence. The identification of an outer membrane protein that is conserved among *E. coli* and some *Shigella* species may also provide potential vaccine and intervention targets to mitigate disease caused by these pathogens. Additionally, this approach may contribute to the discovery of other potential initial adherence factors based on the pursuit of pH-influenced genes that may have a higher probability of involvement in initial adherence or other closely related functions of EHEC. In *S. pneumoniae*, the cell signaling pathway responsible for internalization is also used by other pathogens of diverse natures such as *Staphylococcus aureus, Neisseria meningitidis*, and *Listeria monocytogenes*(Agarwal et al., 2010), suggesting the possibility that this adherence mechanism might be more widespread than previously known.

## Materials and Methods

### Bacterial and mammalian growth and culture

All bacterial strains and plasmids are described in Table 2.

**Table 2.**
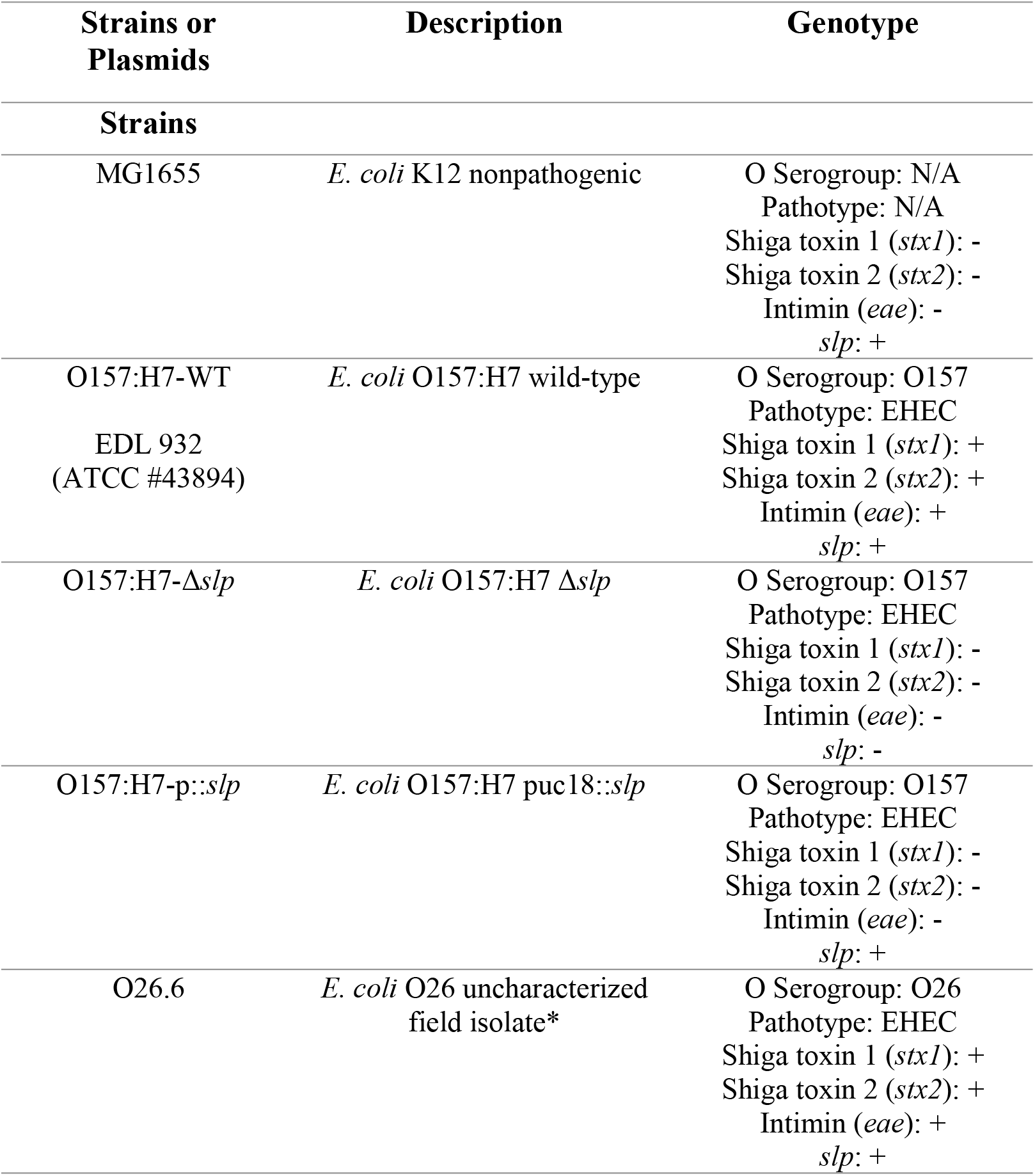
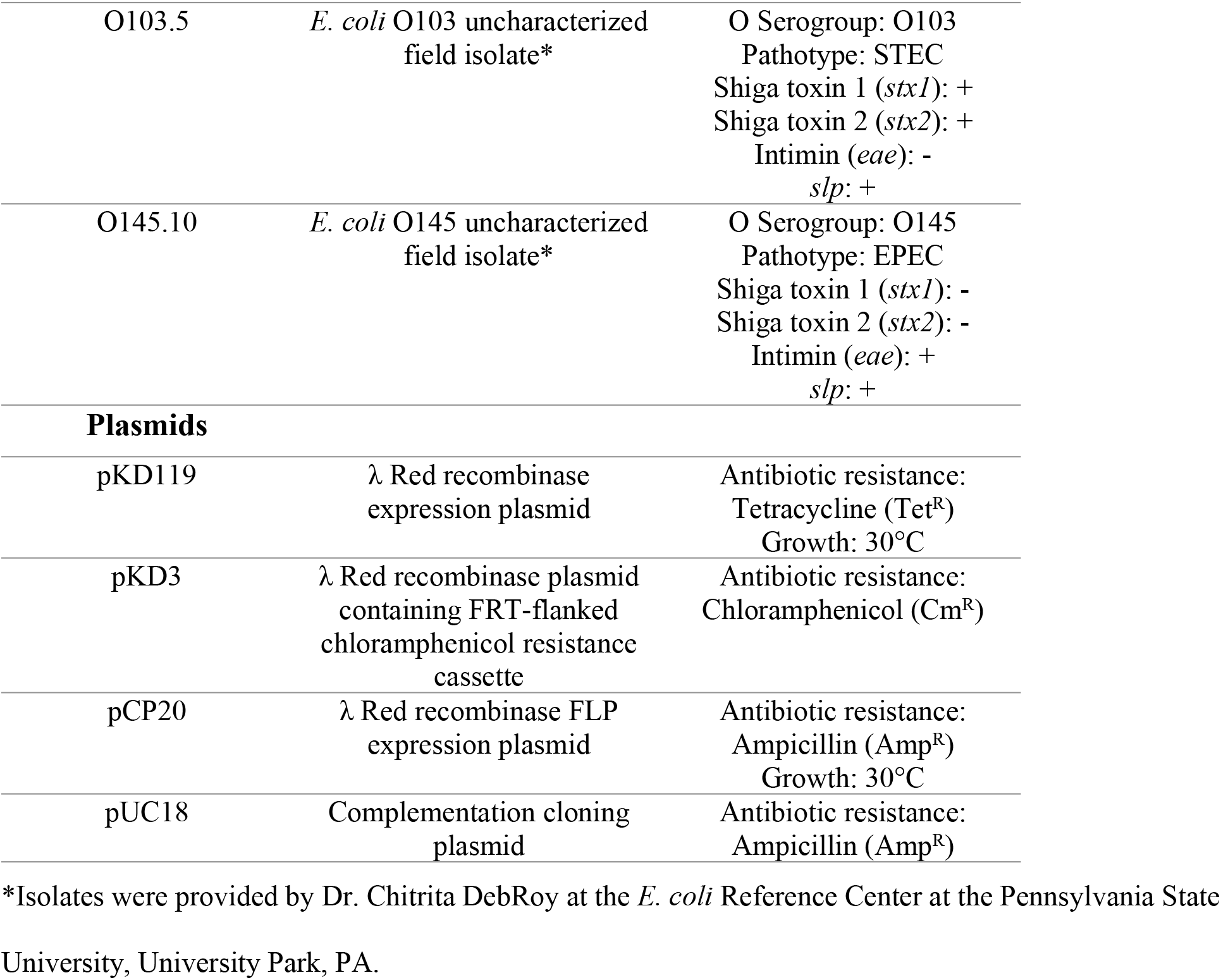
Bacterial strains and plasmids.

| Strains or Plasmids | Description | Genotype |
| --- | --- | --- |
| **Strains** |  |  |
| MG1655 | *E. coli* K12 nonpathogenic | O Serogroup: N/A<br>Pathotype: N/A<br>Shiga toxin 1 (*stx1*): -<br>Shiga toxin 2 (*stx2*): -<br>Intimin (*eae*): -<br>*slp*: + |
| O157:H7-WT<br>EDL 932<br>(ATCC #43894) | *E. coli* O157:H7 wild-type | O Serogroup: O157<br>Pathotype: EHEC<br>Shiga toxin 1 (*stx1*): +<br>Shiga toxin 2 (*stx2*): +<br>Intimin (*eae*): +<br>*slp*: + |
| O157:H7-Δ*slp* | *E. coli* O157:H7 Δ*slp* | O Serogroup: O157<br>Pathotype: EHEC<br>Shiga toxin 1 (*stx1*): -<br>Shiga toxin 2 (*stx2*): -<br>Intimin (*eae*): -<br>*slp*: - |
| O157:H7-p::*slp* | *E. coli* O157:H7 puc18::*slp* | O Serogroup: O157<br>Pathotype: EHEC<br>Shiga toxin 1 (*stx1*): -<br>Shiga toxin 2 (*stx2*): -<br>Intimin (*eae*): -<br>*slp*: + |
| O26.6 | *E. coli* O26 uncharacterized field isolate* | O Serogroup: O26<br>Pathotype: EHEC<br>Shiga toxin 1 (*stx1*): +<br>Shiga toxin 2 (*stx2*): +<br>Intimin (*eae*): +<br>*slp*: + |
| O103.5 | *E. coli* O103 uncharacterized field isolate* | O Serogroup: O103<br>Pathotype: STEC<br>Shiga toxin 1 (*stx1*): +<br>Shiga toxin 2 (*stx2*): +<br>Intimin (*eae*): -<br>*slp*: + |
| O145.10 | *E. coli* O145 uncharacterized field isolate* | O Serogroup: O145<br>Pathotype: EPEC<br>Shiga toxin 1 (*stx1*): -<br>Shiga toxin 2 (*stx2*): -<br>Intimin (*eae*): +<br>*slp*: + |
| **Plasmids** |  |  |
| pKD119 | λ Red recombinase expression plasmid | Antibiotic resistance: Tetracycline (Tet<sup>R</sup>)<br>Growth: 30°C |
| pKD3 | λ Red recombinase plasmid containing FRT-flanked chloramphenicol resistance cassette | Antibiotic resistance: Chloramphenicol (Cm<sup>R</sup>) |
| pCP20 | λ Red recombinase FLP expression plasmid | Antibiotic resistance: Ampicillin (Amp<sup>R</sup>)<br>Growth: 30°C |
| pUC18 | Complementation cloning plasmid | Antibiotic resistance: Ampicillin (Amp<sup>R</sup>) |

\*Isolates were provided by Dr. Chitrita DebRoy at the *E. coli* Reference Center at the Pennsylvania State University, University Park, PA.

All bacterial cultures were inoculated from single colonies into LB broth with appropriate antibiotics when applicable and grown shaking at 37°C with 5% CO_2_ overnight. Caco-2 cells (human colonic adenocarcinoma cells; American Type Culture Collection: ATCC HTB-37) were grown at 37°C with 5% CO_2_, in Eagle’s Minimum Essential Medium (EMEM) (ATCC) plus 20% Fetal Bovine Serum (FBS) (Atlanta Biologicals) unless otherwise indicated. Caco-2 cells were seeded into tissue culture treated six well plates (Corning Life Sciences), containing glass cover slips if used for microscopy, and grown to confluency. pUC18 Complementation cloning plasmid

### Immunofluorescent microscopy

Mammalian cells were grown as described, rinsed once with sterile phosphate buffered saline (PBS) and fresh media applied 2 to 4 hours prior to infection. Bacterial strains were inoculated at an MOI of 20, allowed to equilibrate to 37°C for five minutes, and incubated for 0 to 6 hours. After incubation, cell culture media was removed, and cells were washed gently (by tilting and swirling) with sterile PBS three times to remove unadhered bacteria. After infection, samples were fixed using 10% neutral buffered formalin (Azer Scientific) for ten minutes at room temperature and rinsed with PBS three times. Fluorescent stains were applied as listed below, with two PBS rinses in between each stain; slides were stored in glycerol-based mounting media and away from light. The pIgR protein was stained using an unconjugated PIGR rabbit IgG polyclonal antibody (#PA5–22096, Thermo Fisher Scientific) at a1:750 dilution in PBS for 60 minutes, followed by a DyLight 488-conjugated secondary goat anti-Rabbit IgG (H+L) antibody (#35552, Thermo Fisher Scientific), stained at 1:1,000 dilution in PBS for 60 minutes. *E. coli* were stained using an unconjugated *E. coli* goat IgG polyclonal antibody (#PA1–73032, Thermo Fisher Scientific), stained in 1:1,000 dilution for 60 minutes, followed by a DyLight 594-conjugated secondary donkey anti-goat IgG (H+L) Secondary Antibody (DyLight 594 #SA5–10088, Thermo Fisher Scientific), stained at 1:1,000 dilution in PBS for 60 minutes. Caco-2 cell nuclei were stained using Hoechst 33342 (#62249, Thermo Fisher Scientific), stained at 1:1,000 dilution in PBS for 5 minutes. Slides were washed three times in sterile PBS and mounted using glycerol based mounting media and stored away from light. Immunofluorescent images were obtained using a Keyence BZ-9000 Fluorescence Microscope (Keyence) with 40X objective lenses (Nikon) with filters for DAPI, GFP, Texas-Red, and phase contrast. Images were taken as z-stacks at 0.5 µM intervals. Covariance was measured using ImageJ (U. S. National Institutes of Health) and the JaCOP plugin (64). Statistical analysis was performed using XLSTAT 2017: Data Analysis and Statistical Solution for Microsoft Excel (Addinsoft). P-values were calculated using the mean PCC (R value) for the Student’s T-test and p ≤0.05 was considered significant (65).

### Quantitative adherence

Infection of mammalian cells was done as described. After infection and incubation, samples were collected dispensing 100 µL sterile 10% triton X-100 (Sigma-Aldrich) in PBS and incubated at room temperature for 10 minutes until cells detached from the well plate. A 900 µL volume of sterile PBS was added to rinse the well and samples were collected using a sterile cell scraper for a total sample volume of 1 mL. Samples were resuspended and vortexed vigorously until no visible cell clumps remained, ten-fold serial dilutions were made by adding 100 µL of sample into 900 µL sterile PBS in succession. 100 µL samples of each dilution were plated and grown on LB plates overnight at 37°C before counting. All adherence assays were normalized by calculating colony forming units (CFU) per inoculum to ensure accuracy and performed in triplicate. To calculate adherence, bacterial counts were adjusted by dilution factor, averaged, and the percent adherence (to normalize across samples and experiments) was calculated from the number of adhered bacteria per the number of inoculated bacteria per well. The percent adherence per well was then compared to the maximum adherence count (wild-type count at three hours post-infection) to calculate relative adherence per sample. 
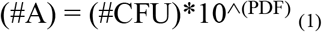

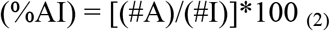

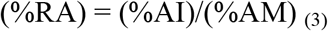

Where (#A) = number of adhered bacteria; (#CFU) = number of colonies counted per plate; (PDF) = plate dilution factor; (#I) = number of inoculated bacteria per well (sample); (%AI) = percent adherence of inoculum (normalization between samples); (%AM) = maximum percent adherence (%AI) of wild-type bacteria (three hour samples); (%RA) = relative percent adherence (values shown in figures). P-values were calculated using the mean %RA for the Student’s T-test and p ≤ 0.05 was considered significant (65).

### Quantitative PCR (qPCR)

Quantitative (real-time) PCR was done using the purified mRNA samples collected as described in Section 2.6 were reverse-transcribed to cDNA using the iScript cDNA Synthesis kit (BioRad). The cDNA samples were then concentrated and cleaned of reverse transcription reaction components using ethanol precipitation: to each sample, 10% (of total reverse transcription reaction volume) volume of 3M sodium acetate pH 5.2 was added; 2.5 volumes of 100% ethanol was added; sample was vortexed thoroughly and incubated at –20°C for a minimum of six hours to form a DNA precipitate. After precipitation, samples were centrifuged (in a standard microcentrifuge) at ˃10,000xg for 30 minutes at 4°C; pellets were rinsed once with ice-cold 70% ethanol; pellets were then air dried and resuspended in water and assessed for purity and concentration using AD 260/280 ratios. Using cDNA as PCR template, 20 µL qPCR reactions were made using 50–500 ng of template cDNA, 10 µL Applied Biosystems SYBR Green PCR Master Mix (Thermo Fisher Scientific), and molecular-grade water up to 20 uL total volume. The qPCR thermal cycling was done using the 7500/7500 Fast Real-Time PCR System (Applied Biosystems) using the default PCR cycling settings (hold stage: 10 minutes at 95°C; cycling stage (40 cycles): 15 seconds at 95°C + 1 minute at 60°C) and the primers listed in Table 3.

**Table 3.**
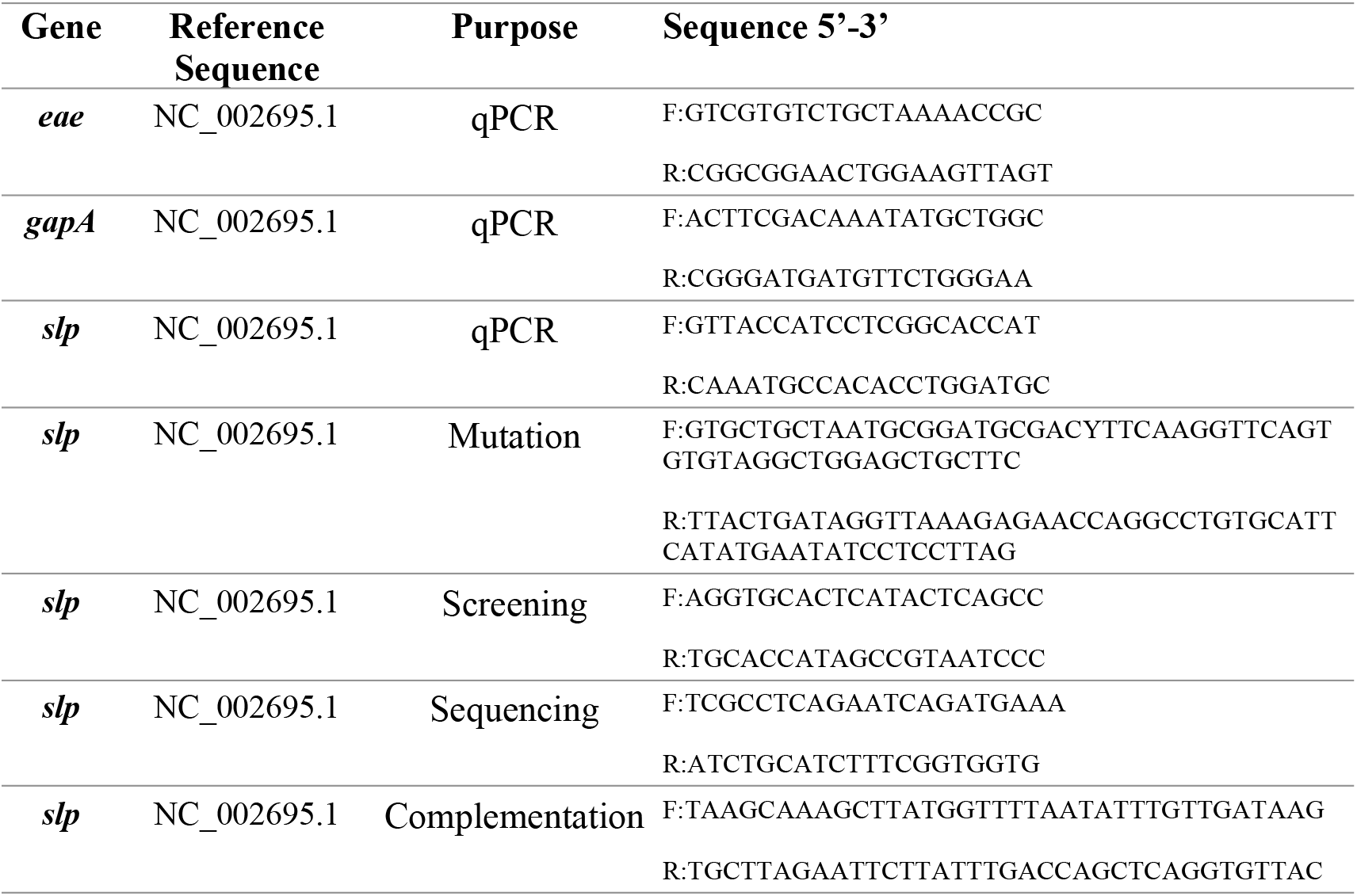
Primer sequences used in this study.

Relative expression was calculated using the relative expression ratio where efficiency (E) was calculated using the slope of a standard curve of ten-fold serial dilutions of DNA template, and fold-change (FC) was calculated using the relative expression ratio as described by Pfaffl (33). 
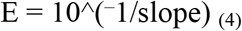

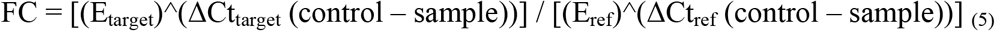
when E = efficiency of target or reference gene, ΔCt = (Ct value of control-Ct value of sample), target = the gene of interest, ref = endogenous control gene, control= untreated sample, and sample = experimental sample (33). A fold-change ≥ 2 was considered significant.

### Gene deletion and complementation

*E. coli* O157:H7 was used to make a genomic deletion lacking *slp* using the lambda Red recombinase system described by Datsenko and Wanner (66). Bacterial strains hosting plasmids were grown with antibiotics according to their respective resistance genes, and plasmids were purified using the Plasmid Midi Kit (Qiagen). PCR, using the mutation primers listed in Table 3, was performed using plasmid pKD3 as a template on an Eppendorf MasterCycler PCR System (Eppendorf). PCR reactions were done in 50 µL volumes: 0.25 µL Taq DNA Polymerase enzyme (5U/µL) (Omega Bio-Tek); 10X PCR buffer (#TQ2100–00, Omega Bio-Tek) at 5 µL; 1.0 µL dNTPs (10mM) (Lucigen); 1.0 µL each of forward and reverse primers (20 µM) (IDT), and water to 50 µL total volume. All PCR cycling conditions were done as follows, with adjusted annealing temperatures noted in Table 2.2: 95°C for 5 minutes; 35 cycles of 95°C for 5 minutes, (Annealing Tm)°C for 30 seconds, 72°C for 3e minutes; followed by a final extension step of 72°C for 5 minutes. PCR products were run in a 1.5% agarose gel in Tris-acetate-EDTA (TAE) buffer at 150 V for 1 to 2 hours to obtain clear band resolution, bands were excised from the gel, and purified using the MinElute Gel Extraction Kit (Qiagen). PCR products were transformed into electrocompetent *E. coli* O157:H7 containing plasmid pKD119, using volumes between 2 and 5 µL at concentrations between 100 and 500 ng/µL (as measured by Nanovue spectrophotometer). Transformed cells were incubated in 1 mL of pre-warmed 37°C SOC medium and incubated at 37°C for 1 hour, then plated on LB containing chloramphenicol. Colonies were selected for antibiotic resistance and screened by PCR. Transformants positive for antibiotic resistance and negative for screening PCR products were grown and made electrocompetent, transformed with plasmid pCP20 to excise the antibiotic resistance cassette, and screened for antibiotic resistance. Deletions were confirmed by sequencing the junction sequences flanking the deletion site, using The Pennsylvania State University’s Genomics Core Facility (University Park, PA). Plasmid complementation of the deletion strain was made using pUC18 and the primers listed in Table 3. PCR was done using *E. coli* O157:H7 genomic DNA as a template; and PCR reactions, PCR cycling conditions, PCR product purification, and transformations were done as described.

### Co-immunoprecipitation (Co-IP) and Protein Identification

The Co-IP assay was done using the Invitrogen Dynabeads Protein A Immunoprecipitation kit (Invitrogen). Bacterial cultures were grown from starter culture diluted into 100 mL LB broth at an OD of 0.05 and grown with shaking to prevent sediment formation for approximately 14 hours at 37°C. Bacterial pellets were collected by centrifugation at 4°C at 4,000xg for 20 minutes and resuspended in 5mL water for sonication. Sonication was done using a hand-held sonicator (Thermo Fisher Scientific) on ice for a total of 20 minutes per sample, or until lysate turbidity was visibly reduced. A 5 mL volume of sonicated bacterial lysate was aliquoted into 1 mL samples. Untreated lysate was not incubated with protein; treated lysate was incubated with 10µg of Recombinant Human Polymeric Immunoglobulin Receptor/PIgR (C-Fc) CI09 protein purchased from Novoprotein (Novoprotein) containing an Fc tag at the C-terminus of the protein. Protein-lysate incubation was done overnight at 4°C for 16–18 hours on a rotator, and proteins collected using the Co-IP kit. Protein A coated beads from the kit were prepared for use and incubated with the lysate for 30 minutes at 4°C on a rotator. Beads were collected using a magnet stand and washed twice with kit wash solution. Beads pellet was then resuspended in 10 µL of kit elution solution and 10 µL of sodium dodecyl sulfate polyacrylamide gel electrophoresis (SDS-PAGE) buffer and incubated at 95°C for 5 minutes before loading into a 12% acrylamide SDS-PAGE gel. The SDS-PAGE gel was run for two hours at 100 V at room temperature and stained with Coomasie blue (BioRad). All protein identifications were done at protein facilities using liquid chromatography and tandem mass spectrometry (LC MS/MS) analysis. A total of three analyses were completed, at The Pennsylvania State Proteomics and Mass Spectrometry Core Facility (1) and the Protein Facility of the Iowa State University Office of Biotechnology (2 and 3). The protein bands were cut from the SDS-PAGE gel and processed at the respective facility. Peptide fragmentation patterns were compared to known databases of proteins of *E. coli* (PSU using SEQUEST and Uniprot; ISU using Mascot or Sequest HT). Raw data was analyzed using Thermo Scientific’s Proteome Discoverer Software.

## Acknowledgments and Disclaimer

Mention of trade names or commercial products in this article is solely for the purpose of providing specific information and does not imply recommendation or endorsement by the US Department of Agriculture (USDA). USDA is an equal opportunity provider and employer.

## References

1. Mead PS, Slutsker L, Dietz V, McCaig LF, Bresee JS, Shapiro C, et al. Food-related illness and death in the United States. Emerg Infect Dis. 1999;5(5):607–25.

2. Hunt JM. Shiga toxin-producing Escherichia coli (STEC). Clin Lab Med. Elsevier Ltd; 2010 Mar [cited 2014 Oct 22];30(1):21–45. Available from: http://www.ncbi.nlm.nih.gov/pubmed/20513540

3. Mohawk KL, O’Brien AD. Mouse Models of *Escherichia coli* O157:H7 Infection and Shiga Toxin Injection. J Biomed Biotechnol. 2011 Jan;2011:1–17. Available from: http://www.hindawi.com/journals/bmri/2011/258185/

4. Rangel JM, Sparling PH, Crowe C, Griffin PM, Swerdlow DL. O157:H7 Outbreaks in the US 1982–2002. 2005;11(4).

5. Schmidt MA. LEEways: tales of EPEC, ATEC and EHEC. Cell Microbiol. 2010 Dec [cited 2014 Oct 31];12(11):1544–52. Available from: http://www.ncbi.nlm.nih.gov/pubmed/20716205

6. Farfan MJ, Torres AG. Molecular mechanisms that mediate colonization of Shiga toxin-producing Escherichia coli strains. Infect Immun. 2012 Mar [cited 2014 Oct 22];80(3):903–13. Available from: http://www.pubmedcentral.nih.gov/articlerender.fcgi?artid=3294676&tool=pmcentrez&rendertype=abstract

7. Bardiau M, Szalo M, Mainil JG. Initial adherence of EPEC EHEC and VTEC to host cells. Vet Res. 2010 [cited 2014 Oct 22];41(5):57. Available from: http://www.pubmedcentral.nih.gov/articlerender.fcgi?artid=2881418&tool=pmcentrez&rendertype=abstract

8. Torres AG, Giron J a, Perna NT, Blattner FR Avelino-flores F, Kaper B, et al. Identification and Characterization of lpfABCC ′DE, a Fimbrial Operon of Enterohemorrhagic Escherichia coli. 2002;70(10):5416–27.

9. Cordonnier C, Etienne-Mesmin L, Etienne-Mesmin L, Thévenot J, Rougeron A, Rénier S, Chassaing B, et al. Enterohemorrhagic Escherichia coli pathogenesis: Role of Long polar fimbriae in Peyer’s patches interactions. Sci Rep. Nature Publishing Group; 2017;7(October 2016):1–14. Available from: http://dx.doi.org/10.1038/srep44655

10. Torres AG, Kanack KJ, Tutt CB, Popov V, Kaper JB. Characterization of the second long polar (LP) fimbriae of Escherichia coli O157:H7 and distribution of LP fimbriae in other pathogenic E. coli strains. FEMS Microbiol Lett. 2004;238(2):333–44.

11. Badea L, Doughty S, Nicholls L, Sloan J, Robins-Browne RM, Hartland EL. Contribution of Efa1/LifA to the adherence of enteropathogenic Escherichia coli to epithelial cells. Microb Pathog. 2003;34(5):205–15.

12. Sharma VK, Kudva IT, Bearson BL, Stasko JA. Contributions of EspA filaments and curli fimbriae in cellular adherence and biofilm formation of enterohemorrhagic Escherichia coli O157:H7. PLoS One. 2016;11(2):1–23.

13. Nicholls L, Grant TH, Robins-browne RM. Identification of a novel genetic locus that is required for in vitro adhesion of a clinical isolate of enterohaemorrhagic Escherichia coli to epithelial cells. Mol Microbiol. 2000;35:275–88. Available from: internal-pdf://106.61.211.232/Nicholls-2000-Identification of a novel geneti.pdf%5Cnhttp://www.ncbi.nlm.nih.gov/pubmed/10652089 http://onlinelibrary.wiley.com/store/10.1046/j.1365-2958.2000.01690.x/asset/j.1365-2958.2000.01690.x.pdf?v=1&t=i06rjn25&s=8f04d

14. Tatsuno I, Horie M, Abe H, Makino K, Shinagawa H, Taguchi H. toxB Gene on pO157 of Enterohemorrhagic Escherichia coliO157 : H7 Is Required for Full Epithelial Cell Adherence Phenotype. Infect Immun. 2001;69(11):6660–9.

15. Sharma VK, Bearson SMD, Bearson BL. Evaluation of the effects of sdiA, a luxR homologue, on adherence and motility of Escherichia coli O157:H7. Microbiology. 2010;156(5):1303–12.

16. Giron J a, Torres a G, Freer E, Kaper JB. The flagella of enteropathogenic Echerichia coli mediate adherence to epithelial cells. Mol Micro. 2002;44:361–79.

17. Hammerschmidt S, Tillig MP, Wolff S, Vaerman J-P, Chhatwal GS. Species-specific binding of human secretory component to SpsA protein of Streptococcus pneumoniae via a hexapeptide motif. Mol Microbiol. 2002 Jan 18;36(3):726–36. Available from: http://doi.wiley.com/10.1046/j.1365-2958.2000.01897.x

18. Mostov KE, Blobel G. A transmembrane precursor of secretory component. J Biol Chem. 1982;257(19):11816–21.

19. Johansen F-E, Kaetzel CS. Regulation of the polymeric immunoglobulin receptor and IgA transport: new advances in environmental factors that stimulate pIgR expression and its role in mucosal immunity. Mucosal Immunol. Nature Publishing Group; 2011 Nov [cited 2014 Oct 20];4(6):598–602. Available from: http://www.pubmedcentral.nih.gov/articlerender.fcgi?artid=3196803&tool=pmcentrez&rendertype=abstract

20. Fernandez MI, Pedron T, Tournebize R, Olivo-Marin JC, Sansonetti PJ, Phalipon A. Anti-inflammatory role for intracellular dimeric immunoglobulin A by neutralization of lipopolysaccharide in epithelial cells. Immunity. 2003;18(6):739–49.

21. Asano M, Komiyama K. Polymeric immunoglobulin receptor. J Oral Sci. 2011;53(2):147–56. Available from: http://joi.jlc.jst.go.jp/JST.JSTAGE/josnusd/53.147?from=CrossRef

22. Kaetzel CS. The polymeric immunoglobulin receptor: bridging innate and adaptive immune responses at mucosal surfaces. Immunol Rev. 2005 Aug;206:83–99. Available from: http://www.ncbi.nlm.nih.gov/pubmed/16048543

23. Gan YJ, Chodosh J, Morgan A, Sixbey JW. Epithelial cell polarization is a determinant in the infectious outcome of immunoglobulin A-mediated entry by Epstein-Barr virus. J Virol. 1997;71(1):519–26. Available from: http://www.pubmedcentral.nih.gov/articlerender.fcgi?artid=191081&tool=pmcentrez&rendertype=abstract

24. Lu L, Lamm ME, Li H, Corthesy B, Zhang J-R. The human polymeric immunoglobulin receptor binds to Streptococcus pneumoniae via domains 3 and 4. J Biol Chem. 2003 Nov 28 [cited 2014 Oct 22];278(48):48178–87. Available from: http://www.ncbi.nlm.nih.gov/pubmed/13679368

25. Dave S, Carmicle S, Hammerschmidt S, Pangburn MK, McDaniel LS. Dual roles of PspC, a surface protein of Streptococcus pneumoniae, in binding human secretory IgA and factor H. J Immunol. 2004;173(1):471–7.

26. Kouki A, Haataja S, Loimaranta V, Pulliainen AT, Nilsson UJ, Finne J. Identification of a novel streptococcal adhesin P (SadP) protein recognizing galactosyl-??1–4-galactose-containing glycoconjugates: Convergent evolution of bacterial pathogens to binding of the same host receptor. J Biol Chem. 2011;286(45):38854–64.

27. Li J, Hovde CJ. Expression profiles of bovine genes in the rectoanal junction mucosa during colonization with Escherichia coli O157:H7. Appl Environ Microbiol. 2007 Apr [cited 2014 Oct 22];73(7):2380–5. Available from: http://www.pubmedcentral.nih.gov/articlerender.fcgi?artid=1855659&tool=pmcentrez&rendertype=abstract

28. Abe H, Tatsuno I, Tobe T, Okutani A, Sasakawa C. Bicarbonate Ion Stimulates the Expression of Locus of Enterocyte Effacement-Encoded Genes in Enterohemorrhagic Escherichia coli O157: H7 Bicarbonate Ion Stimulates the Expression of Locus of Enterocyte Effacement-Encoded Genes in Enterohemorrhagic Escher. 2002;70(7):3500–9.

29. Alsharif G, Ahmad S, Islam MS, Shah R, Busby SJ, Krachler AM. Host attachment and fluid shear are integrated into a mechanical signal regulating virulence in Escherichia coli O157:H7. Proc Natl Acad Sci U S A [Internet]. 2015;112(17):5503–8. Available from: http://www.pubmedcentral.nih.gov/articlerender.fcgi?artid=4418854&tool=pmcentrez&rendertype=abstract

30. Crane JK, Byrd IW, Boedeker EC. Virulence inhibition by zinc in shiga-toxigenic Escherichia coli. Infect Immun. 2011 Apr [cited 2014 Oct 22];79(4):1696–705. Available from: http://www.pubmedcentral.nih.gov/articlerender.fcgi?artid=3067541&tool=pmcentrez&rendertype=abstract

31. Yang B, Feng L, Wang F, Wang L. Enterohemorrhagic Escherichia coli senses low biotin status in the large intestine for colonization and infection. Nat Commun. Nature Publishing Group; 2015;6:6592. Available from: http://www.nature.com/doifinder/10.1038/ncomms7592

32. Yona-Nadler C. Integration host factor (IHF) mediates repression of flagella in enteropathogenic and enterohaemorrhagic Escherichia coli. Microbiology. 2003 Apr 1 [cited 2014 Oct 23];149(4):877–84. Available from: http://mic.sgmjournals.org/cgi/doi/10.1099/mic.0.25970-0

33. Pfaffl MW. A new mathematical model for relative quantification in real-time RT-PCR. Nucleic Acids Res [Internet]. 2001;29(9):e45. Available from: http://www.ncbi.nlm.nih.gov/pubmed/11328886

34. Alexander DM, St John AC. Characterization of the carbon starvation-inducible and stationary phase-inducible gene sip encoding an outer membrane lipoprotein in Escherichia coi. 1994;11:1059–71.

35. Melton-celsa A, Mohawk K, Teel L, Brien AO. Pathogenesis of Shiga-Toxin Producing Escherichia coli. Curr Top Microbiol Immunol. 2012;357(September 2011):67–103.

36. Kalita A, Hu J, Torres AG. Recent advances in adherence and invasion of pathogenic Escherichia coli. Curr Opin Infect Dis [Internet]. 2014;27(5):459–64. Available from: http://www.pubmedcentral.nih.gov/articlerender.fcgi?artid=4169667&tool=pmcentrez&rendertype=abstract

37. Szalo IM, Goffaux F, Pirson V, Piérard D, Ball H, Mainil J. Presence in bovine enteropathogenic (EPEC) and enterohaemorrhagic (EHEC) Escherichia coli of genes encoding for putative adhesins of human EHEC strains. Res Microbiol. 2002;153(10):653–8.

38. Mahajan A, Currie CG, Mackie S, Tree J, McAteer S, McKendrick I, et al. An investigation of the expression and adhesin function of H7 flagella in the interaction of Escherichia coli O157: H7 with bovine intestinal epithelium. Cell Microbiol. 2009;11(1):121–37. Available from: http://www.ncbi.nlm.nih.gov/pubmed/19016776

39. Erdem AL, Avelino F, Xicohtencatl-Cortes J, Girón JA. Host protein binding and adhesive properties of H6 and H7 flagella of attaching and effacing Escherichia coli. J Bacteriol. 2007;189(20):7426–35.

40. Croxen M a, Law RJ, Scholz R, Keeney KM, Wlodarska M, Finlay BB. Recent advances in understanding enteric pathogenic Escherichia coli. Clin Microbiol Rev. 2013 Oct;26(4):822–80.

41. Schneeman T a., Bruno MEC, Schjerven H, Johansen F-EF-EF-EEF-E, Chady L, Kaetzel CS, et al. Regulation of the polymeric Ig receptor by signaling through TLRs 3 and 4: linking innate and adaptive immune responses. J Immunol. Nature Publishing Group; 2005 Jan 13 [cited 2014 Jul 10];175(1):376–84. Available from: http://www.jimmunol.org/cgi/doi/10.4049/jimmunol.175.1.376

42. Bruno MEC, Frantz a L, Rogier EW, Johansen F-E, Kaetzel CS. Regulation of the polymeric immunoglobulin receptor by the classical and alternative NF-κB pathways in intestinal epithelial cells. Mucosal Immunol. Nature Publishing Group; 2011 Jul [cited 2014 Oct 8];4(4):468–78. Available from: http://www.pubmedcentral.nih.gov/articlerender.fcgi?artid=3125104&tool=pmcentre&rendertype=abstract

43. Karpman D, Ståhl A-L. Enterohemorrhagic Escherichia coli Pathogenesis and the Host Response. Microbiol Spectr. 2014;2(5):1–15.

44. Phalipon A, Corthésy B. Novel functions of the polymeric Ig receptor: well beyond transport of immunoglobulins. Trends Immunol. 2003 [cited 2014 Oct 25];24(2):55–8. Available from: http://www.sciencedirect.com/science/article/pii/S1471490602000315

45. Cardone MH, Smith BL, Mennitt PA, Mochly-Rosen D, Silver RB, Mostov KE. Signal transduction by the polymeric immunoglobulin receptor suggests a role in regulation of receptor transcytosis. J Cell Biol. 1996;133(5):997–1005.

46. Price GP, John ACS. Purification and analysis of expression of the stationary phase-inducible Slp lipoprotein in Escherichia coli: Role of the Mar system. FEMS Microbiol Lett. 2000;193(1):51–6.

47. Kovacs-Simon A, Titball RW, Michell SL. Lipoproteins of bacterial pathogens. Infect Immun. 2011;79(2):548–61.

48. Carter MQ, Louie JW, Fagerquist CK, Sultan O, Miller WG, Mandrell RE. Evolutionary silence of the acid chaperone protein HdeB in enterohemorrhagic Escherichia coli O157: H7. Appl Environ Microbiol. 2012;78(4):1004–14.

49. Tramonti A, De Canio M, De Biase D. GadX/GadW-dependent regulation of the Escherichia coli acid fitness island: Transcriptional control at the gadY-gadW divergent promoters and identification of four novel 42 bp GadX/GadW-specific binding sites. Mol Microbiol. 2008;70(4):965–82.

50. Hommais F, Krin E, Coppée JY, Lacroix C, Yeramian E, Danchin A, et al. GadE (YhiE): A novel activator involved in the response to acid environment in Escherichia coli. Microbiology. 2004;150(1):61–72.

51. Zhao B, Houry W a. Acid stress response in enteropathogenic gammaproteobacteria: an aptitude for survival. Biochem Cell Biol. 2010;88(2):301–14.

52. Mates AK, Sayed AK, Foster JW. Products of the Escherichia coli acid fitness island attenuate metabolite stress at extremely low pH and mediate a cell density-dependent acid resistance. J Bacteriol. 2007;189(7):2759–68.

53. Wu VCH, Qiu X, de los Reyes BG, Lin CS, Pan Y. Application of cranberry concentrate (Vaccinium macrocarpon) to control Escherichia coli O157:H7 in ground beef and its antimicrobial mechanism related to the downregulated slp, hdeA and cfa. Food Microbiol. Elsevier Ltd; 2009;26(1):32–8. Available from: http://dx.doi.org/10.1016/j.fm.2008.07.014

54. Schembri MA, Kjaergaard K, Klemm P. Global gene expression in Escherichia coli biofilms. Mol Microbiol. 2003;48(1):253–67.

55. Barnett Foster D. Modulation of the enterohemorrhagic E. coli virulence program through the human gastrointestinal tract. Virulence. 2013;4(4):315–23. Available from: http://www.tandfonline.com/doi/full/10.4161/viru.24318#.VZ0VHUvlfnc%5Cnhttp://www.tandfonline.com/doi/abs/10.4161/viru.24318

56. Masuda N, Church GM. Regulatory network of acid resistance genes in Escherichia coli. Mol Microbiol. 2003;48(3):699–712.

57. Morgan JK, Carroll RK, Harro CM, Vendura KW, Shaw LN, Riordan JT. Global regulator of virulence A (GrvA) coordinates expression of discrete pathogenic mechanisms in enterohemorrhagic Escherichia coli through interactions with GadW-GadE. J Bacteriol. 2015;198(3):394–409.

58. Foster JW. Escherichia coli acid resistance: tales of an amateur acidophile. Nat Rev Microbiol. 2004;2(11):898–907. Available from: http://dx.doi.org/10.1038/nrmicro1021

59. House B, Kus J V, Prayitno N, Mair R, Que L, Chingcuanco F, et al. Acid-stress-induced changes in enterohaemorrhagic Escherichia coli O157: H7 virulence. Microbiology. 2009 Sep [cited 2014 Oct 9];155(Pt 9):2907–18. Available from: http://www.ncbi.nlm.nih.gov/pubmed/19497950

60. Callaway T, Carr M. Diet, Escherichia coli O157: H7, and cattle: a review after 10 years. Curr issues. 2009 [cited 2014 Oct 25];1(979):67–80. Available from: http://www.horizonpress.com/cimb/v/v11/67.pdf

61. Naylor SW, Nart P, Sales J, Flockhart A, Gally DL, Low JC. Impact of the direct application of therapeutic agents to the terminal recta of experimentally colonized calves on Escherichia coli O157:H7 shedding. Appl Environ Microbiol. 2007 Mar [cited 2014 Oct 22];73(5):1493–500. Available from: http://www.pubmedcentral.nih.gov/articlerender.fcgi?artid=1828765&tool=pmcentrez&re

62. Arthur TM, Brichta-Harhay DM, Bosilevac JM, Kalchayanand N, Shackelford SD, Wheeler TL, et al. Super shedding of Escherichia coli O157:H7 by cattle and the impact on beef carcass contamination. Meat Sci. Elsevier B.V.; 2010 Sep [cited 2014 Oct 23];86(1):32–7. Available from: http://www.ncbi.nlm.nih.gov/pubmed/20627603

63. van Diemen PM, Dziva F, Abu-Median A, Wallis TS, van den Bosch H, Dougan G, et al. Subunit vaccines based on intimin and Efa-1 polypeptides induce humoral immunity in cattle but do not protect against intestinal colonisation by enterohaemorrhagic Escherichia coli O157:H7 or O26:H-. Vet Immunol Immunopathol. 2007 Mar 15 [cited 2014 Oct 22];116(1–2):47–58. Available from: http://www.pubmedcentral.nih.gov/articlerender.fcgi?artid=2656997&tool=pmcentrez&rendertype=abstract

64. Datsenko K a, Wanner BL. One-step inactivation of chromosomal genes in Escherichia coli K-12 using PCR products. Proc Natl Acad Sci USA. 2000 Jun 6;97(12):6640–5. Available from: http://www.pubmedcentral.nih.gov/articlerender.fcgi?artid=18686&tool=pmcentrez&rendertype=abstract

65. Mcdonald JH, Dunn KW. Statistical tests for measures of colocalization in biological microscopy. J Microsc. 2013;252(3):295–302.

